# Epidermal Growth Factor potentiates EGFR(Y992/1173)-mediated therapeutic response of triple negative breast cancer cells to cold atmospheric plasma-activated medium

**DOI:** 10.1101/2023.05.30.542966

**Authors:** Peiyu Wang, Renwu Zhou, Rusen Zhou, Lihui Yu, Liqian Zhao, Wenshao Li, Jinyong Lin, Aleksandra Rajapakse, Chia-Hwa Lee, Frank B. Furnari, Antony W. Burgess, Jennifer H. Gunter, Gang Liu, Kostya (Ken) Ostrikov, Derek J Richard, Fiona Simpson, Xiaofeng Dai, Erik W. Thompson

## Abstract

Cold atmospheric plasma (CAP) holds promise as a cancer-specific treatment that selectively kills basal-like breast cancer cells. We used CAP-activated media (PAM) to capture the multi-modal chemical species of CAP. Specific antibodies, small molecule inhibitors and CRISPR/Cas9 gene-editing approaches showed an essential role for receptor tyrosine kinases, especially epidermal growth factor (EGF) receptor, in mediating triple negative breast cancer (TNBC) cell responses to PAM. EGF also dramatically enhanced the sensitivity and specificity of PAM against TNBC cells. Site-specific phospho-EGFR analysis, signal transduction inhibitors and reconstitution of EGFR-depleted cells with EGFR-mutants confirmed the role of phospho-tyrosines 992/1173 and phospholipase C gamma signaling in upregulating levels of reactive oxygen species above the apoptotic threshold. EGF-triggered EGFR activation enhanced the sensitivity and selectivity of PAM effects on TNBC cells, such that a strategy based on the synergism of CAP and EGF therapy may provide new opportunities to improve the clinical management of TNBC.

## INTRODUCTION

Breast cancer (BC) is one of the most prevalent and malignant cancers in women, and affects men at a lower rate (1/100^th^). Among various breast cancer subtypes, triple negative breast cancers (TNBCs) are known for their poor clinical outcome and dearth of effective targeted therapies with acceptable toxicity profiles, due to the absence of druggable targets seen in other breast cancer subtypes ^1,2^. There is a need to explore novel therapeutic modalities that, in combination with current therapies, can help improve clinical outcomes and reduce side effects, especially in TNBC. One such potential therapy is cold atmospheric plasma (CAP), which has proven selectivity against many types of cancer cells ^3^ and represents an emerging oncotherapeutic approach offering a new opportunity to effectively manage a wide range of cancers.

CAP, the plasma generated at room temperature and atmospheric pressure, has shown great potential for biomedical applications ^4–8^, due to a favorable combination of reactive physical and chemical species, such as UV radiation, electrons, free radicals, ions and excited molecules ^9^. More recently, plasma activated medium (PAM) produced by exposure of aqueous medium to CAP^10^, was also found to inhibit cancer cells, sometimes as effectively as the direct CAP treatment ^11^, increasing the scope and flexibility of plasma-based therapeutics ^12^. We have previously reported the remarkable anti-cancer capacity of PAM against TNBCs ^13,14^, and here identify a critical role for epidermal growth factor receptor (EGFR), which is highly expressed in TNBC ^15–17^, and its response to PAM. Increasing evidence has associated high EGFR with immune resistance ^18^ and with high PD-L1 expression ^19^. Synergies between PAM and EGF were identified in this project, which explored the underlying biochemical mechanisms triggered in TNBCs upon treatment with PAM or PAM and EGF together. Combined treatment of TNBC cells with EGF and PAM caused remarkably potent and selective cell killing that is contradictive of conventional anti-cancer approaches using EGFR inhibitors, through cooperative increases in intracellular levels of reactive oxygen and nitrogen species (RONS).

## RESULTS

### EGFR signaling contributes to the selectivity of PAM against TNBC cells

TNBC cells were sensitive, in a dose-dependent manner, to PAM treatment. Treatment with 70% 10PAM (i.e., PAM that was prepared by treating cell culture medium with CAP for 10 min, **Figure 1a**) induced approximately 40% apoptosis in MDA-MB-231 and SUM-159PT cells, and more than 70% apoptosis was seen when cells were treated with 100% 10PAM (**Figure 1d**). Optical emission spectra (OES) analysis was used to identify the excited molecular species generated at the argon plasma-liquid interface (**Figure 1b**). Bands corresponding to the second positive system of N_2_ were observed in the range of 300-400 nm. OH, the rotational band (A^2^Σ+ → X^2^Π) at 308.8 nm, strong atomic argon emission lines in the range of 690-800 nm, and a relatively weak oxygen peak at 777 nm were also clearly observed to originate from the interactions of argon plasma, ambient air and water. The long-lived constituent species of PAM, including H_2_O_2_, NO_3_^−^, NO_2_^−^, and O_3_ were generated by treating 1.5 mL of DMEM with plasma for 10 min, and their concentrations determined to be 50 mg L^−1^, 1000 mg L^−1^, 20 mg L^−1^, and 0.9 mg L^−1^, respectively (**Figure 1c**). The effects of PAM on the viability of BC cell lines were also time-dependent. Across all the monitored durations of 10PAM exposure, cell apoptosis became evident at 6 h, and reached the highest significance at 24 h, where 70% 10PAM-induced apoptosis was highest in TNBC cell lines (**Figure 1e** and **Figure S1a-S1c**).

**Figure 1.**
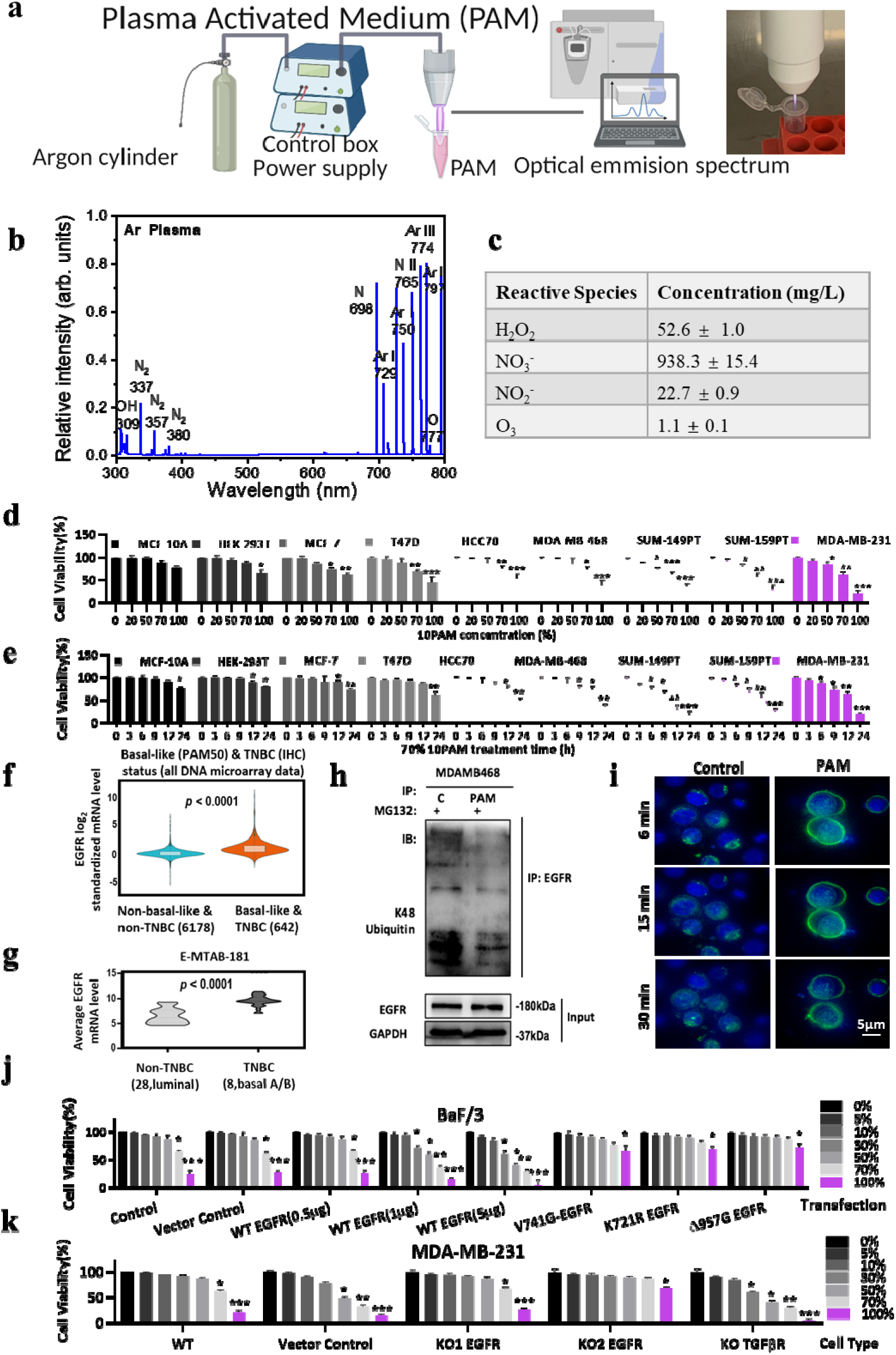
Experimental evidence showing the essential role of EGFR signaling in the selectivity of PAM against TNBC cells *in vitro*. **a** Schematic graph of the CAP device set up. Model presentation of the CAPs generation and an image of kINPen 09 for CAP generation. **b** Optical emission spectra for argon plasma generated using kINPen 09. **c** Concentrations of activated species in 10PAM (10 min-CAP activated PAM). **d** Effects of 12 h treatment with control (C) or 20%, 50%, 70% and 100% 10PAM as indicated on viability (% of C; mean +/− SEM) of breast cancer cell lines (pink shading of TBNC lines, HCC70, MDA-MB-468, SUM-149PT, SUM159-PT, MDA-MB-231 reflects levels of responses to PAM). Cells were plated in 96-well plates at 5,000 cells per well for 24 h and serum free starvation before treatments. **f** Effects of different time durations (0 h, 3 h, 6 h, 9 h, 12 h and 24 h) of 70% 10PAM on viability of different cell lines as per (**d**). **f** EGFR expression according to basal-like PAM50 molecular profile and TNBC immunohistochemistry (IHC) staining status in the METABRIC cohort from bc-GenExMiner v4.5. **g** EGFR gene expression among 28 luminal and 28 basal A/B breast cancer cell lines in the E-MTAB-181 dataset. **h** EGFR ubiquitination levels in MDA-MB-468 cells with or without PAM treatment. **i** MDA-MB-468 cells were plated in an 8-well plate at 2×10^5^ cells per well for 24 h. After 30min pre-treatment with 0% PAM (Control) or 50% PAM, 10 ng mL^−1^ GFP-488 labelled EGF was added to the medium. The images were taken by Delta Vision Ultra High-Resolution Microscope at 6 min, 15 min and 30 min after adding EGF. **j** Viabilities (% of 0% PAM control; mean +/− SEM) of Ba/F3 cells transfected with WT or mutant EGFR in response to PAM, as indicated. **k** Viabilities of MDA-MB-231 cells (% of C; mean +/− SEM) with and without Crispr/Cas-9 suppression of EGFR or TGFβR, in response to PAM of different doses.

EGFR is known to respond actively to oxidative stress ^20^, and given the reactive oxygen and nitrogen species (RONS) enriched in PAM ^21^ and the selectivity of PAM against TNBC cells (**Figure 1d-e**, pink shade), we examined EGFR expression levels across different breast cancer subtypes using public datasets. EGFR expression is higher in basal-like breast cancer cells ^16,17^, most of which are TNBC according to the Prediction Analysis of Microarray 50 (PAM50) and IHC status in the METABRIC dataset as assessed with bc-GenExMiner v4.5 (**Figure 1f**, p<0.0001), and at the transcriptional level from the E-MTAB-181 dataset ^22^ as assessed with ArrayExpress ^23^ (**Figure 1g**, p<0.0001). EGFR signaling is controlled by endocytosis after ligand activation ^24^. Accordingly, we examined the effects of PAM on EGFR ubiquitination and EGFR trafficking in MDA-MB-468, SUM-159PT and MDA-MB-231 TNBC cells. EGFR ubiquitination was substantially reduced upon 50% 10PAM treatment (**Figure 1h, S1d**) and its diffuse plasma membrane staining became more concentrated on the plasma membrane after PAM treatment, particularly at points where two cell membranes meet / overlap (**Figure S1e**) in MDA-MB-468. A peri-nuclear enrichment was also especially evident in the SUM-159PT and MDA-MB-231 cells (**Figure S1f**). This may reflect delayed or altered exocytic pathways as anti-EGFR steady state imaging shows the entire cellular EGFR while EGF-Alexa 488 time course show the kinetics of endocytosis. Using green-fluorescence (Alexa 488)-labelled EGF and high-resolution microscopy, we observed that whilst EGF was endocytosed in control (untreated) cells, 50% 10PAM-treated cells retained EGF on the cell surface even at 30 min post 50% 10PAM treatment (**Figure 1i**). Thus, PAM caused enhanced EGFR stability and amplified EGFR-mediated cytoplasm signaling through EGF membrane retention in TNBC cells.

The murine, interleukin (IL)-3 dependent, BaF/3 hematopoietic cell line, which lacks endogenous EGFR ^25^, was used to explore the role of EGFR in PAM cell responsiveness using EGFR transfection. Although both 70% and 100% 10PAM treatments induced BaF/3 cell death in the control (untreated) and mock-transfected BaF/3 cells, the more EGFR transfected into the BaF/3 cell line (from 0.5 µg to 5 µg), the more the cells were killed by the PAM treatment (**Figure 1j**). BaF/3 cells transfected with 5 µg EGFR were sensitive to 30% 10PAM, with 72.48% cell viability. By transfecting BaF/3 with plasmids encoding the wildtype EGFR and different domain mutants, only cells transfected with the wildtype EGFR expression were more sensitive to 10PAM treatment; there was no increased effect of 10PAM on cell killing when cells were transfected with mutants encoding abrogated EGFR kinase activation (V741G-EGFR, K721R EGFR, Δ957G EGFR). These data demonstrated the importance of EGFR signaling in facilitating its enhancement of PAM efficacy against cancer cells.

To explore this role further, we silenced EGFR in MDA-MB-231 human breast cancer cells using CRISPR/Cas9 gene editing, and used the KO2 cell line, in which EGFR was undetectable by Western blot (**Figure S1g**). The KO1 cell line, where EGFR was reduced but not silenced, was used as a negative control. Increased EGFR expression was observed in the KO TGFβR cell line, which was also used as a positive control. The KO2 EGFR cell line was the most resistant to PAM treatment, with 76% cell viability observed at 100% PAM, while the KO TGFβR was the most sensitive to PAM, with only 14% of cells surviving after being treated with 100% 10PAM (**Figure 1k**).

EGFR-specific tyrosine kinase inhibitors erlotinib (Tarceva), gefitinib (Iressa), afatinib (Gilotrif), and the EGFR-specific monoclonal antibodies panitumumab (PmAb, Vectibix) and cetuximab (CmAb, Erbitux), were used to further explore the role of EGFR signaling in killing of TNBC cells by 100% 10PAM for 24h (**Figure S2a**). The EGFR inhibitors significantly reduced the effects of PAM on TNBC, cells but not in the luminal MCF-7 cell line, consistent with the results of **Figure 1d-e**. This further confirmed an important role of EGFR signaling in cell response to the PAM treatment.

A mouse experiment undertaken using the SUM-159PT TNBC cell line (**Figure 2a**) confirmed the PAM-induced repression of tumour weight (0.640±0.855g to 0.064±0.095g; p=0.0113), and showed that the EGFR antagonist afatinib could significantly (p=0.0327) weaken the anti-cancer effect of PAM; tumor weight was increased from 0.064±0.095g to 0.224±0.272g (**Figure 2b-c**) without any vital organ damage (**Figure 2c**). The pilot *in vivo* study showed similar results (**Figure S2b**). Immunohistochemistry staining of EGFR(Tyr992), EGFR, Ki67 and caspase-3 in SUM-159PT tumors from these mice (**Figure 2e**) consolidated the role of CAP in activating EGFR(Tyr992) (**Figure 6c**), enhancing EGFR stability (**Figure 1h, S1d**), blocking EGFR nuclear translocation (**Figure 1i, S2e-f**), inhibiting tumor cell growth (**Figure 1d-e**), and triggering cancer cell apoptosis (**Figure 3f**).

**Figure 2.**
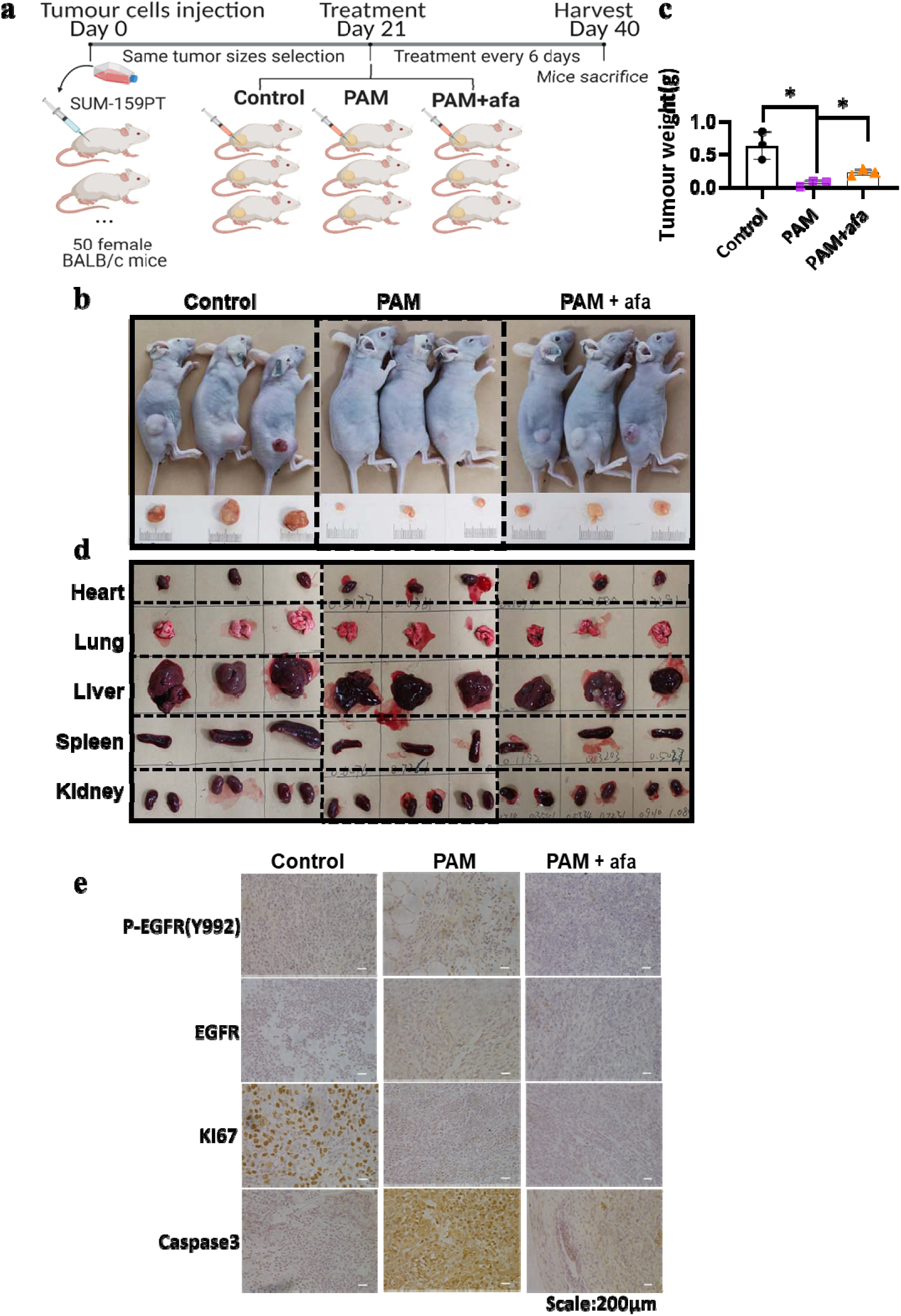
Experimental evidence showing the essential role of EGFR signaling in the selectivity of PAM against TNBC cells *in vivo*. **a** Schematic illustration of the mouse trial testing PAM treatment of SUM-159PT cells in the absence or presence of EGFR inhibitor afatinib (afa). **b** Results from the *in vivo* showing the effects of EGFR inhibition with afatinib (afa) on 50%PAM-treated SUM-159PT tumors. **c** Quantification on the tumor size in the mice. The tumor weight (mean +/− SEM) of control group mice, PAM only treatment and PAM + afatinib treatment. **d** Images of heart, lung, liver, spleen and kidney of mice carrying SUM-159PT tumors in the control, 50%PAM-treated and 50%PAM-treated plus EGFR inhibition with afa. **e** Immunohistochemistry staining results of EGFR(Tyr992), EGFR, Ki67, Caspase 3 on SUM-159PT carrying mice samples in the control, CAP-treated, and CAP-treated plus afatinib groups.

### EGF creates synergies with PAM towards enhanced efficacy against TNBC cells

When EGF (10 ng/mL) was used together with 50% 10PAM for 12h, the potency of 10PAM against TNBC cells was strongly enhanced. While the viability of non-TNBC cell lines was not reduced as much as TNBC, a significant reduction in cell viability was observed for TNBC cells (43.2% for HCC70 with p=0.0008; 73.20% for MDA-MB-468, 58.12% for SUM-159PT and 76.17% for MDA-MB-231 with p<0.0001), whereas less reduction was observed in luminal breast cancer cell lines, T47D and MCF-7 (45.05% for T47D with p=0.001, 17.1% for MCF-7 with p=0.0398) cells (**Figure 3a**).

**Figure 3.**
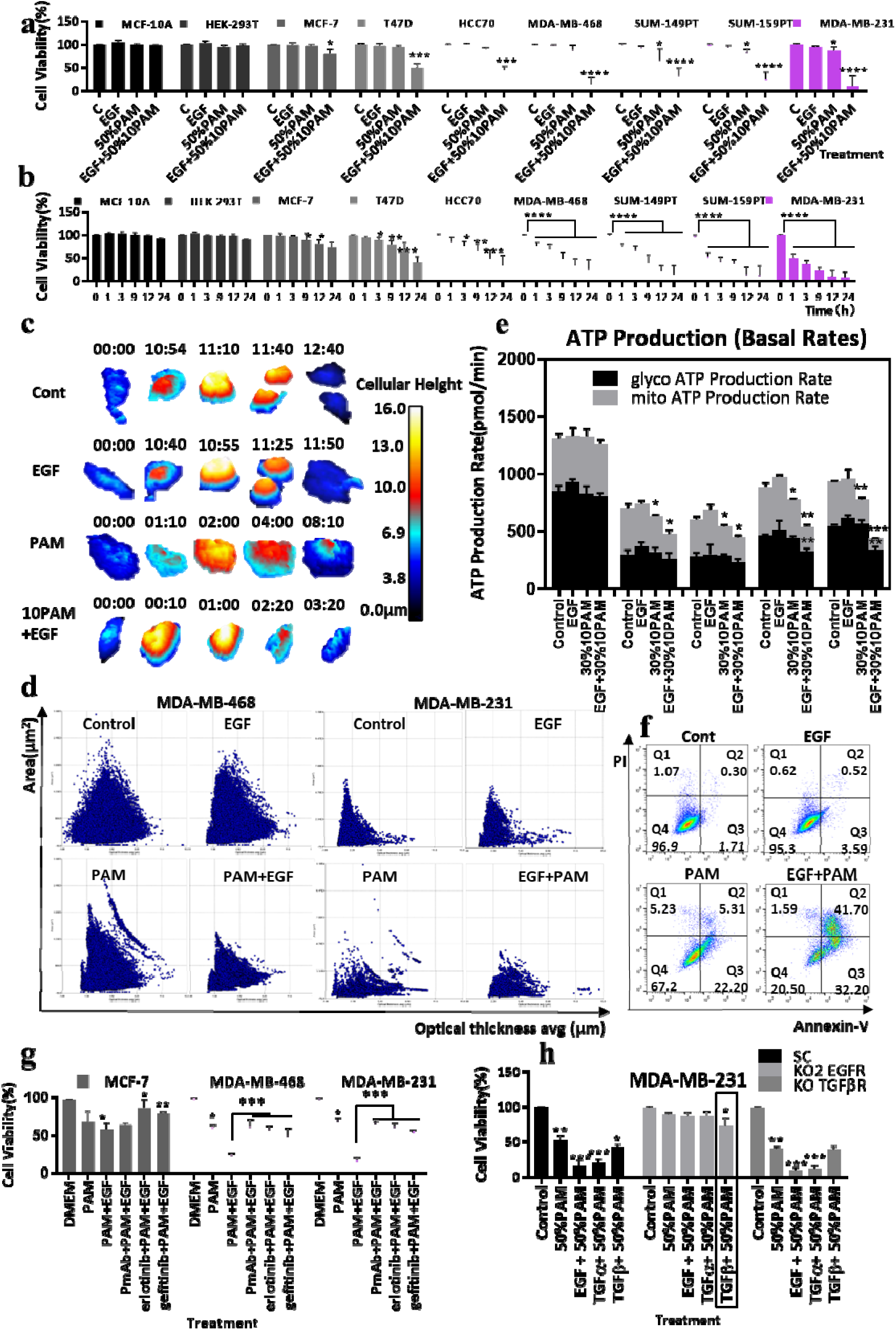
EGF enhances the selectivity of PAM against TNBC cells. **a** Effects of treatment with 50% 10PAM, 10 ng mL^−1^ EGF or both for 12h on viability (% of control (C); mean +/− SEM; N=3) of breast cancer cell lines as indicated (pink shading of TBNC lines: HCC70, MDA-MB-468, SUM-149PT, SUM159-PT, MDA-MB-231). **b** Effects of EGF, 50%PAM or EGF+50%PAM treatments for 12h as in (**a**), for different time durations as indicated, on cell viability (% of control (C); mean +/− SEM; N=3). **c** Representative HoloMonitor images (taken every 5 min, 3 days in total) of MDA-MB-231 cell examples treated with EGF, 50%PAM, or EGF+50%PAM. The time points of cell division (Control and EGF group) and apoptosis (PAM and PAM+EGF group) have been selected for display. **d** Cell area (µm^2^) and optical thickness avg (µm) of MDA-MB-468 and MDA-MB-231 cells measured by the Holomonitor Hstudio after EGF, PAM or EGF+PAM treatments as in (**a**) above, for 3 days, versus Control in MDA-MB-468 and MDA-MB-231 cultures. **e** ATP production rates by MCF-10A, MCF-7, T47D, MDA-MB-468 and MDA-MB-231 (from left to right) upon Control, EGF, PAM or EGF+PAM treatments, for 30 min. OCR and ECAR were measured by Seahorse XF Analyzer to determine rates of glycolysis (glycol) or OXPHOS (mito). **f** Cell apoptosis determined by PI and Annexin V analysis of MDA-MB-231 cells after 50%PAM, EGF or 50%PAM+EGF treatments (as per (**a**) above) for 1 h. **g** Viabilities (% of control (DMEM); mean +/− SEM; N=3) of MCF-7, MDA-MB-468 and MDA-MB-231 cells plated for 24h and pre-treated with 50 ng mL^−1^ PmAb, 1 nM erlotinib and 17 nM gefitinib for 1 h before being treated with 50%PAM, 50%PAM+EGF, or 50%PAM+EGF plus EGFR inhibitors as above for 12 h. **h** Viabilities (% of Control; mean +/− SEM; N=3) of MDA-MB-231 variants (SC, KO2 EGFR, KO TGFβR) treated with 50% PAM, 10 ng mL^−1^ EGF+ 50% PAM, 10 ng mL^−1^ TGFα+ 50% PAM, 10 ng mL1 TGFβ+ 50% PAM for 12 h.

EGF also significantly (p<0.0001) enhanced the sensitivity of BC cells to 50%PAM treatment. While a minimum of 12 h was required for TNBC cells to be responsive to PAM treatment, only 1-3 h was required to trigger equivalent cell apoptosis when EGF and PAM were used together (**Figure 3b**).

Morphology of MDA-MB-231 cells as viewed in the Holomonitor changed from a spindle-like shape to a small and roundish shape after the combined treatment of 50%PAM and EGF for 3 h (**Figure 3c**). While MDA-MB-231 cell division in the control group was observed at 12h, PAM-induced apoptosis was evident at 8 h and cell apoptosis was initiated at 3 h when cells were exposed to combined EGF and 50% 10PAM treatment. Under the control conditions, the correlation between cell area (µm^2^) and optical thickness avg (µm) as measured by Holomonitor Hstudio was evident in video imaging (**Figure 3d** and **Figure S3a**-**b**, **S4a-b**). When plotted against each other, the control group showed an equilateral-like triangular shape, indicating that untreated MDA-MB-468 cells had equivalent area and height, with a roundish ball shape. After 30 min EGF treatment, cells became much taller but had a smaller area than the control group (**Figure S4c-d**). The single 50% 10PAM treatment caused cells to adopt a taller and broader shape as compared with the other groups, suggesting necrosis ^26,27^. PAM synergizes with EGF in causing altered cell morphology in terms of area, height, volume and motility speed, and the number of cells decreased dramatically (**Figure S4e-h**).

Rates of ATP production by mitochondrial respiration and glycolysis were assessed simultaneously using an Agilent Seahorse XFe96 Analyzer (**Figure 3e**). Pre-treatment of cells with 30% PAM for 30 min significantly decreased mitochondrial ATP generation in MCF-7 (p=0.0446), T47D (p=0.0361), MDA-MB-468 (p=0.0369) and MDA-MB-231 (p=0.0029) cells, but not MCF-10A cells (p=0.681), which is consistent with our observation that MCF-10A cells were unresponsive to PAM treatment. Although 30% PAM treatment for 30 min did not reduce cell viability, it was sufficient to influence cells’ energy production. Furthermore, combined treatment inhibited approximately 60% of the oxidative phosphorylation (OXPHOS) (p=0.0049 for MDA-MB-468, p=0.0096 for MDA-MB-231) and part of the glycolysis of MDA-MB-468 and MDA-MB-231 (p=0.0039 for MDA-MB-468, p=0.0009 for MDA-MB-231). Suppressed glycolysis may contribute to the increased sensitivity of TNBCs to PAM treatment when PAM was combined with EGF (**Figure S5a-c**).

The proportion of apoptotic and necrotic cells after 50%PAM and/or EGF treatment for one-hour was determined by Annexin V and PI double staining. Quadrants Q1, Q2, Q3 and Q4 denote necrotic cell (Annexin V−/PI+), late apoptotic cell (Annexin V+/PI+), early apoptotic cell (Annexin V+/PI−) and viable cell (Annexin V−/PI−) regions, respectively (**Figure 3f**). EGF treatment did not cause any significant change as compared with the control. 50%PAM treatment increased the percentages of both necrosis and apoptosis, while combined 50%PAM + EGF treatment boosted the apoptotic proportion. Among cells receiving combined PAM and EGF exposure, 74% (32%+42%) underwent apoptosis and only 2% underwent necrosis (p<0.0001), which was consistent with Holomonitor observations.

Cell viability was significantly reduced by 70%-80% in TNBC cells (MDA-MB-231, MDA-MB-468) when cells were treated with combined 50%PAM and EGF for 3 h, whereas the viability retention was rescued when EGFR inhibitors (PmAb, erlotinib, geftinib) were added (**Figure 3g**). The 50%PAM + EGF combination did not inhibit cell viability in the MCF-7 cell line, supporting the selectivity of PAM against TNBC cells and the role of EGF triggered EGFR signaling in mediating such an efficacy. These results indicated that both antibodies and tyrosine kinase inhibitors could inhibit the synergies between EGF and PAM in killing of EGFR-overexpressing cancer cells.

Gene editing to remove EGFR from MDA-MB-231 cells significantly reduced the proportion of cells killed by 10PAM (p=0.001) (**Figure 3h**). The same results were obtained when 12h EGF and 50%PAM were applied together (p=0.0003), or when cells were treated with a combination of PAM and TGFα (p=0.0001), another EGFR ligand (**Figure 3h**). Other growth factors such as TGFβ and FGFs also showed a synergistic effect with 50%PAM, depending on the cell line and the growth factor (**Figure S5g**). TGFβ (p=0.0175) also enhanced the reduction of cell viability when combined with PAM in the absence of EGFR in the KO2 cells (**Figure 3h**), indicating that TGFβR also enhances the efficacy of PAM rather than EGFR crosstalk. Besides, the titration of 10 PAM (**Figure S5d**) and EGF (**Figure S5e**) suggested the determinant role of PAM rather than EGF in their synergies in cell killing, with 0.01 ng/mL EGF sufficient to synergize with PAM in triggering cell apoptosis. Treatment order also played a role here, with EGF pre-treatment more efficient than PAM pre-treatment in reducing cancer cell viability (**Figure S5f**).

### EGFR activation by PAM + EGF enhances cellular ROS level that trigger cell apoptosis

Since reactive species overload has been implicated in the selective killing of cancer cells by PAM ^9^, we tested the effects of PAM and/or EGF on intracellular ROS. Using the same panel of cell lines as in **Figure 1d-e**, we found that treatment with either EGF or 50% PAM for 12 h had minimal effects on the ROS level, whereas the combined treatment of 50% PAM and EGF selectively enhanced the cellular ROS level in TNBC cells (**Figure 4a**). Combined 50% PAM and EGF treatment for 2 h did not alter cellular ROS level of the normal cells (MCF-10A and HEK-293T), increased the ROS level of luminal cell lines (MCF-7, T47D and HCC70), and substantially enhanced the ROS level of TNBC cell lines (MDA-MB-436, SUM-159PT, MDA-MB-231) (**Figure 4b**). In MDA-MB-231 cells, the ROS level reached a peak of 520% of control at 1 h and remained at a high level even after 6 h. Cellular ROS levels increased with PAM treatment dose regardless of the cell type (**Figure S6a-c**), and the resting state intracellular ROS levels increased in the order of normal, luminal and TNBC cell lines (**Figure S6d**). A tight correlation (R^2^=0.7502) was found between the constitutive ROS level of the cell lines and cell viability responses to 100% PAM treatment (**Figure 4c**), and the strength of this correlation increased (R^2^= 0.9247) when 30% PAM+EGF was used (**Figure 4d**).

**Figure 4.**
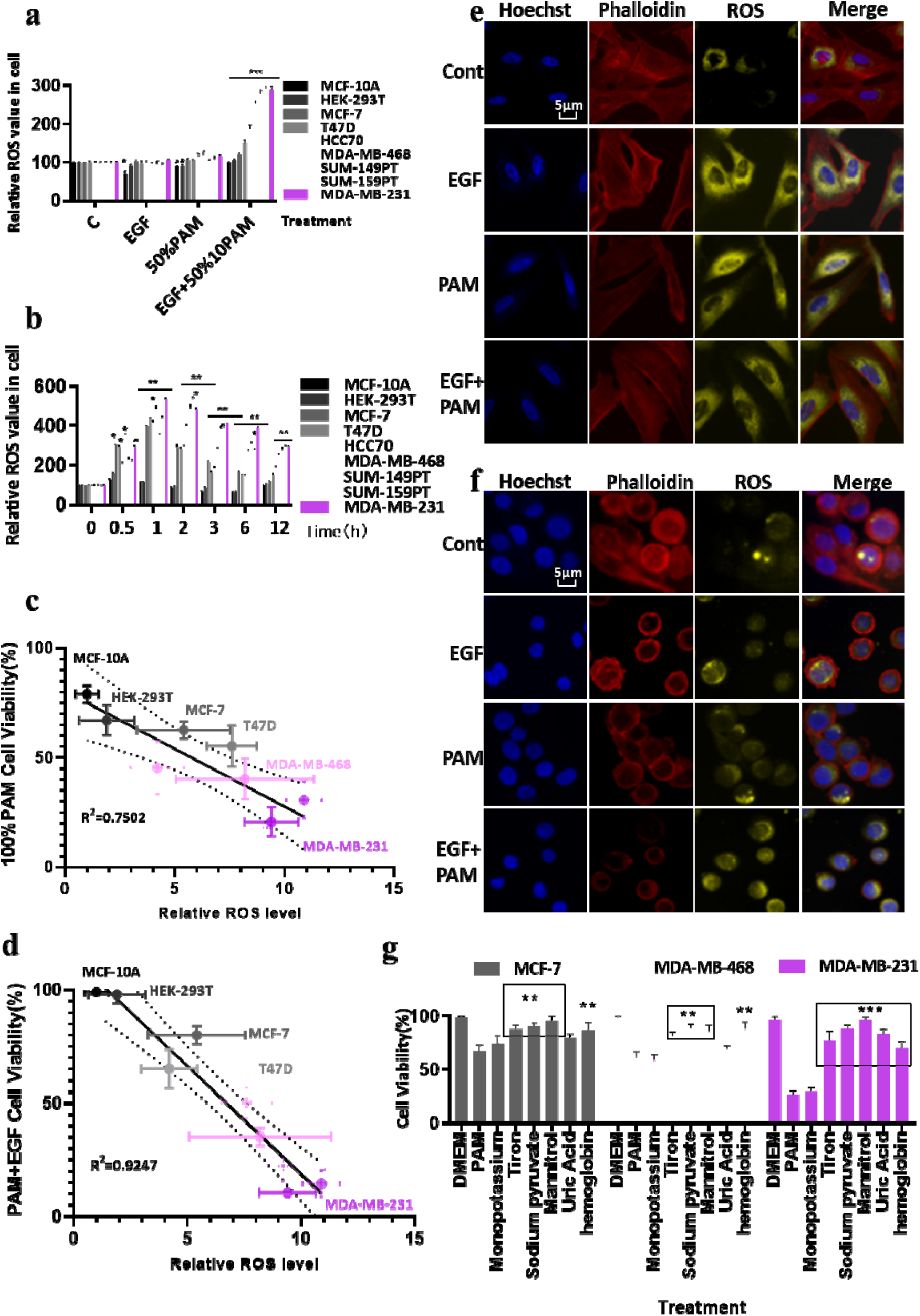
EGF+PAM enhances cellular ROS level that triggers cell apoptosis. **a** Relative ROS levels (% of Control; mean +/− SEM; N=3) of different cell lines (pink shading of TBNC: HCC70, MDA-MB-468, SUM-149PT, SUM159-PT, MDA-MB-231) upon 10 ng mL^−1^ EGF, 50% 10PAM, 50% 10PAM+10 ng mL^−1^ EGF treatment treatments for 12h as indicated. and then incubated with ROS detection kit for 30 min. **b** Relative ROS level of different cell lines upon treatment with 50% 10PAM+10 ng mL^−1^ EGF treatment as per (**a**), for different durations as indicated. **c** Plot showing the linear correlation between ROS levels (Arbitrary Units) of the different cell lines and cell viability (% of Control) in response to 100% PAM treatment for 12 h. R^2^= 0.7502. **d** Plot showing the linear correlation between ROS level of different cell lines and cell viability as per (**b**) in response to 50% 10PAM+10ng mL^−1^ EGF treatment for 12 h. **e** Fluorescence imaging showing cellular ROS (incubated with ROS detection kit for 30 min), actin architecture (phalloidin stain), and nuclei (Hoechst 33342) in MCF-10A and **f** MDA-MB-468 cells treated with EGF, PAM, EGF+PAM for 3 h from (**b**). **g** Viabilities (% of Control; mean +/− SEM; N=3) of MCF-7, MDA-MB-468 and MDA-MB-231 cells pre-treated for 1h with control or scavengers of different ROS components (200 mM mannitol, 100 μM uric acid, 20 mM tiron, 20 μM hemoglobin, 10 mM sodium pyruvate and 1 mM monopotassium were used to quench hydroxyl radical, ozone, superoxide anion, nitric oxide, H_2_O_2_, and e^−^, respectively), and then treated with 50% 10PAM.

ROS levels were also assessed by immunofluorescence in MCF-10A (**Figure 4e**) and MDA-MB-468 (**Figure 4f**) cells. While ROS levels in malignant MDA-MB-468 cells were tripled by combined treatment (50%PAM + EGF) as compared with PAM treatment alone, this did not cause any significant alteration in MCF-10A cells. Cell membrane-associated actin (indicated by phalloidin red) and cytoplasmic distribution of ROS remained consistent after PAM or combined PAM+EGF treatment in MCF10A cells. In MDA-MB-468 cells, cell membrane-associated actin was reduced by PAM and abrogated by the combined PAM+EGF treatment as compared with the control (no treatment), and ROS accumulated in cell nuclei (**Figure 4f** and **S6e**). Collectively, these results suggested a role of ROS in PAM-triggered cell apoptosis and, in particular, in the combined PAM+EGF treatment in BC cells.

By quenching each active component of CAP and examining cell viabilities, we found that all ROS quenchers, except for monopotassium that abrogates the effect of solvated electrons (e^−^) ^28^, rescued cell viability from 50%PAM or 50%PAM+EGF treatment in MDA-MB-231 cells (**Figure 4g, Figure S6f**). This suggested that hydroxyl radical (OH; mannitrol), superoxide anion (O_2_^−^; tiron), nitric oxide (NO; hemoglobin), (H_2_O_2_; sodium pyruvate), as well as reactive species generated from their interactions, all played roles in mediating the selectivity of PAM against TNBC cells.

### The EGFR (Tyr992/1173) axis is essential in mediating the synergistic effect of PAM and EGF

Given our accumulated evidence on the synergistic role of EGF and PAM in killing TNBC cells, we were motivated to identify the specific EGFR phosphorylation sites ^29,30^ mediating these effects. MDA-MB-231 cells, which are highly sensitive to both 50%PAM and combined 50%PAM+EGF treatments were chosen for further studies. Based on the configuration map of the multi-EGFR phosphorylation assay (**Figure S7m**), phosphorylation of Tyr992 and Tyr1173 were up-regulated by PAM and combined PAM + EGF treatments, while phosphorylation of Tyr1045, Tyr1068, Tyr1086, Tyr1148, Ser1045/1047 and Ser1070 were down-regulated by these treatments (**Figure 5a-b**). The Tyr845 site remained active even after 3 h serum starvation, suggesting that this could be a constitutive autophosphorylation site in the MDA-MB-231 cell line.

**Figure 5.**
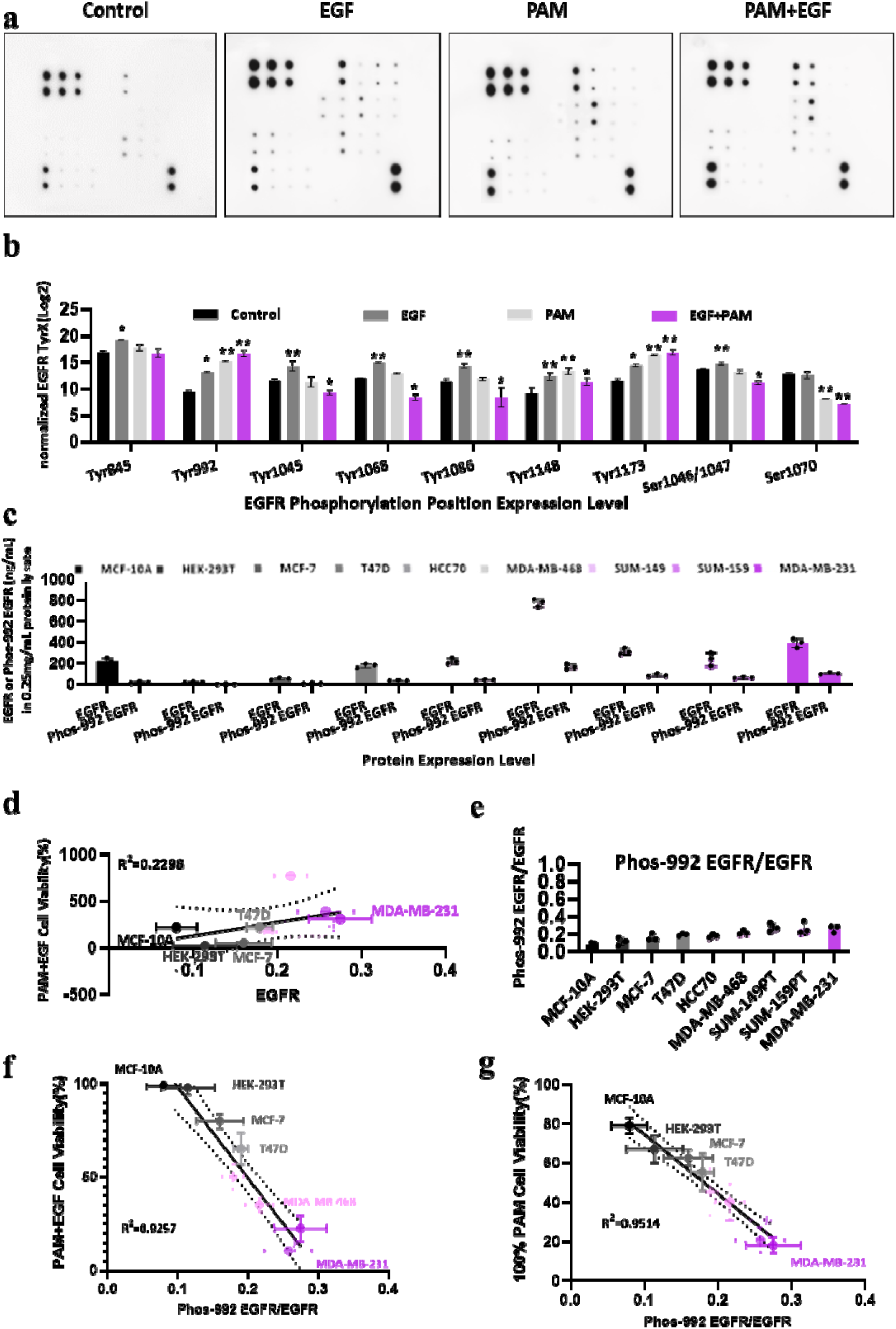
The EGFR (Tyr992/1173) axis is essential in mediating the synergistic effect of PAM and EGF. **a** Multi-site EGFR phosphorylation assay of MDA-MB-231 cells showing EGFR phosphorylation site activation status upon EGF, 50%PAM, EGF+50%PAM treatments. Cells were plated in 6-well-plates at 5×10^5^ per well for 24 h. After 3 h starvation with serum-free medium, the cells were treated with or without EGF (10 ng mL^−1^), 50% 10PAM or PAM+EGF. The protein lysates were collected and normalized by protein assay. Human EGFR Phosphorylation Antibody Array Membranes (17 Targets; ab134005) were used to quantitate the degree of phosphorylation at each site, as per the MATERIALS AND METHODS. **b** Quantification of (**a**). Graph of the quantification data from (**a**) normalised to pan EGFR, as detailed in MATERIALS AND METHODS. Each membrane was scanned and measured by Image Lab 3 times. **c** Levels of EGFR phosphor-Tyr992-EGFR in different cell lines. The nine cell lines were plated in 6-well-plates at 5×10^5^ per well for 24 h. Their protein lysates were collected and normalized to 25 mg/ml by protein assay. The STAR EGFR ELISA Kit was utilized to measure EGFR and levels of Tyr992 phosphorylated-EGFR was measured by EGFR-Y992 ELISA, as per MATERIALS AND METHODS. N=3. **d** Linear correlation between total EGFR levels and viability (% of control) after 50% 10PAM+10 ng mL^−1^ EGF treatment. R^2^=0.2298. **e** Proportion of Tyr992-phosphorylated EGFR (Phos-992 EGFR) to total EGFR in different untreated cell lines. TNBC cell lines shown in pink, with the color intensity representing the Phos-992EGFR/EGFR level. **f** Linear correlation between resting Phospho-992 EGFR/ total EGFR ratio and cell viability (% of control) in response to PAM+EGF treatment. R^2^=0.9257. **g** Linear correlation between Phospho-992 EGFR/ total EGFR and viability in response to 100% PAM. R^2^=0.9514, as per (**f**).

ELISA measurement of total and phospho-Tyr992 EGFR levels in the 9 cell lines (**Figure 5c**) showed that among the TNBC lines, MDA-MB-468 cells had the highest EGFR level followed by the basal B cell lines (SUM-149PT, SUM-159PT and MDA-MB-231) ^31^. Interestingly, the EGFR level in MCF-10A cells was similar to that in T47D and HCC70 cells, while MCF-7 and HEK-293T cells displayed almost no EGFR expression (**Figure 5c**). Although there was not a close relationship between EGFR level and cells’ sensitivity to PAM or PAM+EGF treatments (R^2^ =0.2298, **Figure 5d**), the ratio of phosho-Tyr992 EGFR to total EGFR in untreated cells showed a high correlation with their sensitivities to PAM (R^2^= 0.9514) or PAM+EGF (R^2^= 0.9257) exposure (**Figure 5e-f**).

### Phospholipase C gamma (PLC**γ**, PLCG1) is involved in the synergistic effects of PAM and EGF

To identify the EGFR residues that mediated CAP-triggered EGFR signaling, we transfected EGFR phosphorylation site-mutants into the EGFR-silenced KO2 MDA-MB-231 cell line (**Figure 1k**). The genomic sequencing (**Figure S7a-S7e**) and the multi-EGFR phosphorylation assay (**Figure S7f-S7l**) confirmed the successful construction of these EGFR mutants. When treated with PAM (**Figure 6a**) and 50%PAM+EGF for 12h (**Figure 6b**), cells transfected with the wild type EGFR were sensitive to PAM treatment, consistent with the results of **Figure 1k**. Cells transfected with Tyr1068-, Tyr1086- or Try1148-mutated EGFR were more sensitive to PAM and combined PAM+EGF exposure than cells transfected with the wild type EGFR, presumably because these site-mutated forms of EGFR could not influence the activation of PI3K/AKT survival signaling. Importantly, cells carrying Tyr992- or Tyr1173-mutant EGFR remained resistant to PAM or PAM+EGF treatments, with 100% PAM and PAM+EGF triggering only 30% and 28.7% apoptosis, respectively. Similar results were observed in cell lines transfected with DY5- or DY6-mutated EGFR plasmids, where DY5-mutant included Tyr992, Tyr1068, Tyr1086, Tyr1148 and Tyr1173 and DY6-mutant included all these sites plus Tyr845. The results collectively confirmed our findings of the primary role of Tyr992 and Tyr1173 in mediating PAM or PAM+EGF triggered cell response.

**Figure 6.**
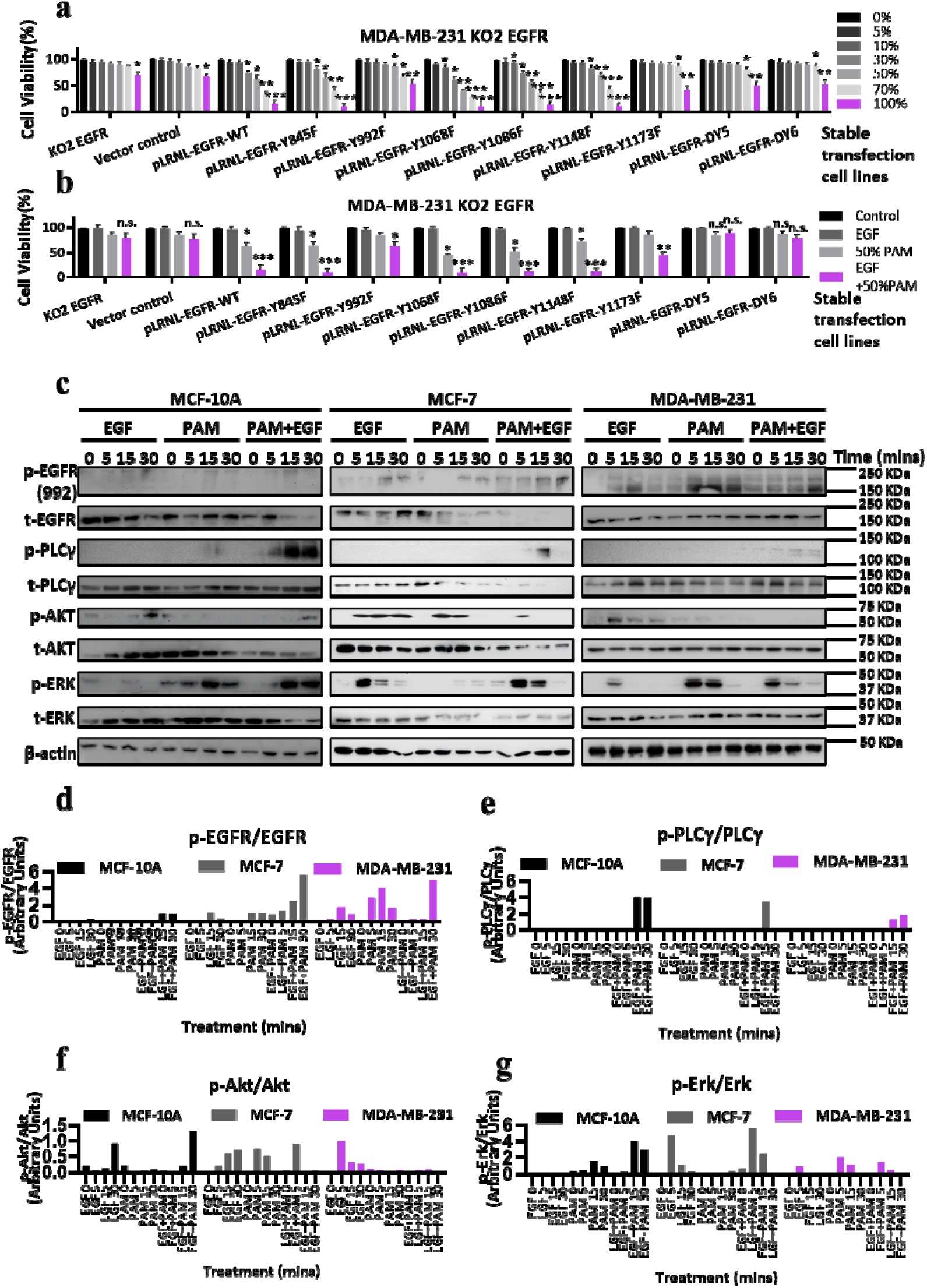
PLCγ is involved in the synergistic effect of PAM and EGF. **a** Viabilities (% of control; 0% PAM) of EGFR-MDA-MB-231 cells (KO2 EGFR, Figure 1n) stably re-transfected by vector only (pLRNL plasmid), pLRNL-EGFR-WT, pLRNL-EGFR-Y845F, pLRNL-EGFR-Y992F, pLRNL-EGFR-Y1068F, pLRNL-EGFR-Y1086F, pLRNL-EGFR-Y1148F, pLRNL-EGFR-Y1173F, pLRNL-EGFR-DY5 or pLRNL-EGFR-DY6 as indicated. These cell lines were plated in 96-well-plates at 5,000 cells per well and treated with 0%, 5%, 10%, 30%, 50%, 70% and 100% 10PAM for 12h followed by cell viability test, with 3 replicates. **b** Viabilities (% of control) of MDA-MB-231 KO2 EGFR cells re-transfected with vector, wildtype EGFR or different EGFR mutations as per (**a**) in response to 10 ng mL^−1^ EGF, 50% 10PAM and EGF+50%PAM for 12h followed by cell viability test, with 3 replicates. **c** Western blots of p-992 EGFR, total EGFR level, p-PLCγ, total PLCγ, p-AKT, total AKT, p-ERK, total ERK in MDA-MB-231, MCF-7 and MCF-10A cells under different treatment conditions. Cells were plated in 12-well-plate at 2.5×10^5^ per well for 24 h, and treated with 10 ng mL^−1^ EGF, 50% 10PAM or EGF+PAM for 0, 5, 15, 30 min after 3 h starvation in serum-free medium. The protein lysates were collected and normalized by protein assay. Quantification is shown for p-992 EGFR/EGFR (**d**), p-PLCγ/ PLCγ (**e**), p-AKT/AKT (**f**) and p-ERK/ERK (**g**). Each sample has 3 replicates.

To identify the signaling downstream of EGFR Tyr992/Tyr1173, we assessed the levels of EGFR, PLCγ, AKT and ERK, and their phosphorylated forms via Western blotting of lysates from MCF-10A (normal), MCF-7 (luminal) and MDA-MB-231 cells (TNBC) (**Figure 6c**). EGFR Tyr992 phosphorylation was highly activated by the combined 50%PAM+EGF treatment in these three cell lines (**Figure 6d**). PLCγ phosphorylation was only detectable after combined 50%PAM+EGF treatment in all three cell lines (**Figure 6e**). PLCγ was activated in MCF-10A and MDA-MB-231 cells when incubated with 50%PAM for 15 to 30 min; PLCγ activation was observed in MCF-7 cells at 15 min 50%PAM + EGF incubation but decreased after 30 min. These results were consistent with the observation that MCF-7 cells were less sensitive to combined treatment than MDA-MB-231 cells. Phosphorylated AKT (p-Akt) was activated by EGF in all three cell lines but was inactivated by 50%PAM and totally inhibited by the combined 50%PAM+EGF treatment in MDA-MB-231 cells (**Figure 6f**). Although it was activated by 50%PAM in MCF-7 cells, it only lasted for 15 min on PAM treatment, and only for 5 min with 50%EGF+PAM treatment. Interestingly, p-AKT was stimulated after 30 min of combined PAM+EGF treatment in MCF-10A, reflecting the survival of normal cells under PAM-imposed stress. ERK activation was upregulated by EGF, PAM and their combined treatment in MDA-MB-231 and MCF-10A cells (**Figure 6g**). Through examining multiple signaling pathways downstream of EGFR using various inhibitors, we identified PLCγ as the most significantly activated target for 50%PAM responses (p=0.0186 for MCF-7, p=0.0006 for MDA-MB-468, p=0.0050 for SUM-159PT and p=0.0042 for MDA-MB-231, **Figure S8**), offering one potential molecular mechanism that drives the selectivity of PAM and its synergy with EGF in killing TNBC cells.

### The EGFR (Tyr992/1173) is necessary for PAM induced ROS level increase and ATP generation reduction

Accumulating results indicated that phosphor-Tyr992/1173 EGFR and PLCγ were essential for CAP-induced apoptosis by increasing enormous ROS generation and decreasing the metabolic rate. We were motivated to test the ROS level and ATP production rate of the EGFR phosphorylation site-mutants. Among these mutants, only cells stably transfected with EGFR-Y992F and Y1173F could downregulate ROS generation response to PAM/PAM+EGF, while cells transfected with EGFR-WT, Y845F, Y1068F, Y1086F and Y1148F dramatically increased the ROS value in cells under 12h 50% or 100%PAM and joint treatments (**Figure 7a-b**). In terms of ATP production, 30 min 10% or 30%PAM treatment or EGF joint treatment could not affect the metabolic rate of KO2 EGFR cells, but obviously reduced both glycolysis and OXPHOS metabolism of EGFR-WT cells and these mutants including Y845F, Y1068F, Y1086F and Y1148F. However, Y992F and Y1173F reversed PAM- or (PAM+EGF)-triggered reduction on glycolysis and OXPHOS (**Figure 7c-d**; **Figure S9a**-**b**). These results indicated the necessity of Tyr992/1173 in endogenous ROS production and ROS-triggered ATP reduction.

**Figure 7.**
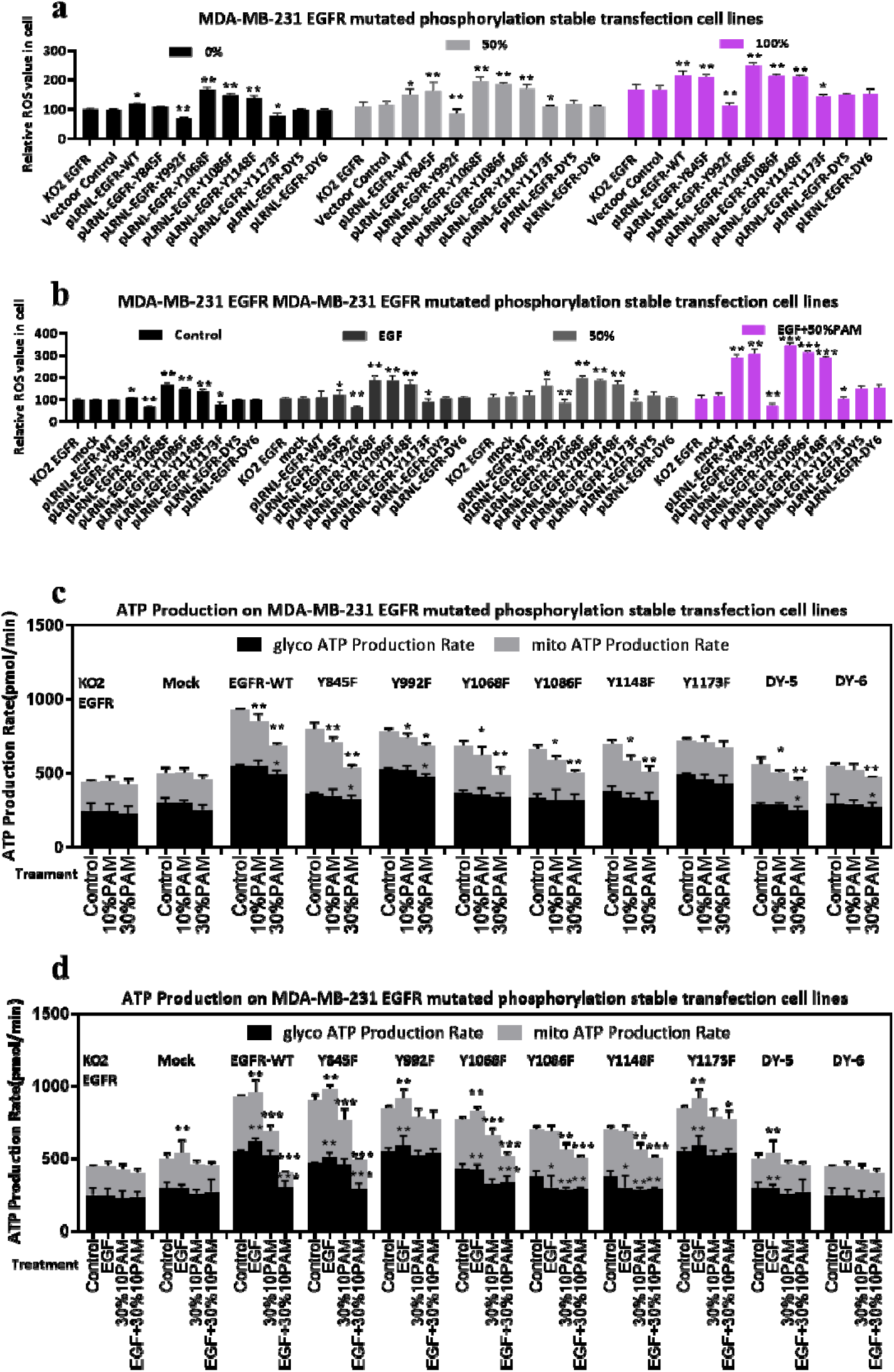
The EGFR (Tyr992/1173) is necessary for PAM-induced ROS increases and reduction in ATP generation. **a** Relative ROS levels (% of KO2 EGFR control+/− SEM, N=3) of MDA-MB-231 KO2 EGFR cells re-transfected with vector, wildtype EGFR or different EGFR mutations upon 0, 50, and 100% PAM treatment at different doses as indicated. The stably transfected cells were seeded in 96-well-plates at 5,000 cells per well for 24 h and treated with or without 50% or 100% PAM for 12h. Cells were then incubated with the ROS detection kit for 30 min and stained by Hoechst 33342 and phalloidin. The cell photos were taken by InCell 6500HS and ROS level was calculated by IN Carta Analysis. **b** Relative ROS levels (% of KO2 EGFR control +/− SEM, N=3) of MDA-MB-231 cells carrying vector control, wildtype EGFR or different EGFR mutations, treated with or without 10 ng mL^−1^ EGF, 50% 10PAM or EGF+PAM for 12 h, after which the ROS levels were tested as in (**a**). **c** ATP production rates in MDA-MB-231 KO2 EGFR cells re-transfected with vector, wildtype EGFR or different EGFR mutations, in response to PAM treatment at different doses. Cells were seeded in Seahorse plates at 20,000 cells per well for 24 h. After 3 h starvation in serum-free medium, cells were treated with or without 10% PAM or 30% PAM for 30min. OCR and ECAR were measured by Seahorse XF Analyzer following the manufacturer’s recommended protocol to determine rates of glycolysis (glycol) or OXPHOS (mito). **d** ATP production rates as in (**c**) above, in MDA-MB-231 KO2 EGFR cells re-transfected with vector, in response to EGF, 30% PAM, EGF+30% PAM treatments.

## DISCUSSION

We show in this study that receptor tyrosine kinases (RTK) ligands such as EGF can dramatically promote PAM-induced apoptosis of TNBC cells, where EGF acts through EGFR-Tyr 992/1173/PLCγ signaling (**Figure 8**) in association with its existing degree of activation (**Figure 5e, 5f**).

**Figure 8.**
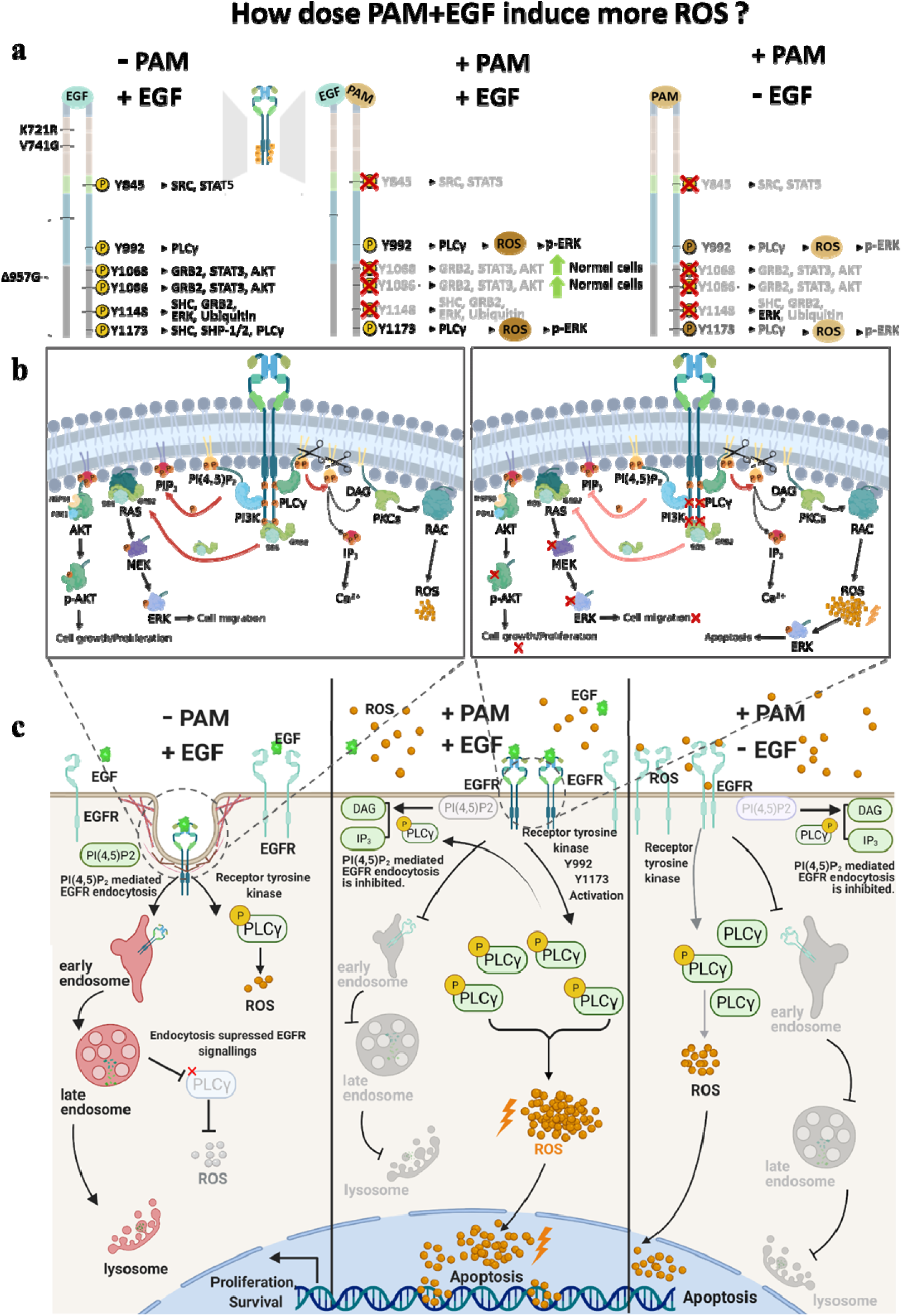
Schematic representation of the mechanism of activation of EGFR by –PAM/+EGF, +PAM/+EGF, or +PAM/-EGF, and the respective downstream signaling. **a** Auto-phosphorylation of EGFR kinase activation sites (K721, V741) that cause the conformational changes to allow autophosphorylation of other sites. When EGF binds to the EGFR ligand binding domain, EGFR undergoes structural changes to form a dimer and the phosphorylation sites are exposed, and activate various downstream signaling pathways; pTyr845 recruits SRC protein and activate STAT5 signaling, pTyr1068 and pTyr1086 recruits GRB2 and phosphorylase STAT3 and AKT. pTyr1148 recruits SHC and GRB2 activating ERK and ubiquitin. Both pTyr 992 and pTyr1173 trigger PLCγ signaling to increase endogenous ROS generation, which in turn leads to apoptosis. The +PAM/+EGF and +PAM/-EGF treatments selectively activate specific phosphorylation sites (Tyr992/1173). While phosphorylation of Tyr845, Tyr1068, Tyr1086 and Tyr1148 is inhibited, phosphorylation of both Tyr992 and Tyr1173 are enhanced, to trigger PLCγ signaling and increase endogenous ROS generation. This is more evident in cancer cells, which tend to already have higher ROS levels, however we found that Tyr1068 and Tyr1086 could still be activated by +PAM/+EGF, +PAM/-EGF in normal cells. **b** PI(4,5)P_2_ mediates three defined signaling events, including (**i**) EGFR endocytosis and stimulation of (**ii**) PLCγ/DAG/PKC/RAC/ROS and (**iii**) PI3K/AKT. EGF-activated EGFR dimer activates these at the same time. As only PLCγ could be activated during the +PAM/+EGF or +PAM/-EGF treatments, the PI(4,5)P_2_ is mainly converted to DAG and IP_3_ to recruit PKCs and RAC, generating ROS, due to the effect of competitive inhibition, rather than PIP_3_. **c** As the ubiquitination and endocytosis of EGFR are inhibited by PAM/EGF+PAM treatment, the phospho-Tyr992/Tyr1173/EGFR/ PLCγ/ROS pathway could be continually activated, unlike the EGF-triggered pathway activation, which will be inhibited in 30 min due to endocytosis and ubiquitination. ROS has also been shown to activate ERK phosphorylation, and this can have secondary effects on other cellular functions such as proliferation and migration. The graph created by Biorender.com (access on 10^th^ June 2021).

We assessed cell apoptosis, viability, cell morphology, and ATP production rate in this study to evaluate the role of EGF in promoting PAM-triggered cell responses. The altered cell morphology from spindle-like to roundish upon combined treatment with PAM and EGF (**Figure 3c**) is likely due to a cell response to EGF stimulation of integrin attachment to the extracellular matrix ^32^. As higher glycolytic and mitochondrial ATP production are associated with higher metabolism and thus higher cell proliferation rate, which is one hallmark of cancers ^33,34^, we included this phenotypic index when assessing the efficacy of PAM, as well as its synergy with EGF, in triggering cell apoptosis. Both glycolytic and mitochondrial ATP production rates were considerably higher in TNBC cells, and were significantly (p<0.0001) reduced on exposure to the combined use of EGF and PAM, suggesting increased apoptosis and effective malignancy control with PAM and, in particular, with the combined use of EGF and PAM. In addition, the ATP production rates of TNBC cells were reduced to a level comparable with the luminal type of cancer cells once treated with joint PAM + EGF treatment, suggesting a possible transition in the metabolic state of TNBC cells from a hormone receptor-irresponsive to a -responsive state.

EGFR activation has been associated with a cell proliferative role, as well as migration, invasion, and epithelial mesenchymal transition ^35–39^ and is necessary for the PAM response, however, it is largely blocked by small molecule inhibitors erlotinib and gefitinib, afatinib, and the antibodies panitumumab and cetuximab. Consistent with this, mutation of the EGFR kinase completely blocked PAM response, even in the absence of EGF, and blocked the response to combined treatment. These observations together suggest that PAM should possibly not be used with EGFR inhibitors such as erlotinib, gefitinib or afatinib in cancer treatment.

Reasonable consensus exists on the role of ROS played in PAM-induced cancer cell cytotoxicity ^3,9^. Indeed, there was a high correlation between the resting ROS level and cell response to PAM, with the highest ROS level observed in the most aggressive TNBC cell lines ^31^, as previously reported ^12^. In contrast, the quasi-normal MCF-10A cell line had very low levels of ROS and showed a very blunted response, and the less aggressive luminal breast cancer cell lines, e.g., MCF-7 and T47D, showed low levels of ROS and reduced cell responses. Consistent with the prevailing dogma, increases in the ROS levels observed after the combined EGF and PAM treatment were also higher in the more responsive TNBC cell lines. Thus, our data is supportive of the positive relationship between cellular ROS levels and PAM responsiveness ^40^.

Although potentially toxic to cells, ROS also play important signaling roles downstream of RTKs such as EGFR and have been implicated in EGFR-mediated tumor progression and therapy resistance ^41^. ROS such as H_2_O_2_ are generated by NADPH-dependent oxidases in response to EGFR stimulation, and have a second messenger function to regulate intracellular signaling cascades by modifying specific cysteine and methionine residues within redox-sensitive protein targets ^42,43^. We focused on the functionality of EGFR(p-Tyr992/Tyr1173) in PAM-mediated cell responses by locating EGFR phosphorylation sites that showed largest differences between PAM and PAM+EGF treatment. MDA-MB-231 cells were selected as the studying system as it exhibited the most obvious synergistic effect and was PAM-sensitive (**Figure 3a**), rendering the identified molecular mechanism extensible to other TNBC cells. Among the EGFR phosphorylation sites we examined, Tyr1045, Tyr1068, Tyr1086, Tyr1148, Ser1046/1047, Ser1070 were also significantly affected by the synergistic treatment with PAM and EGF, in addition to Tyr992 and Tyr1173. However, Tyr992 and Tyr1173 were the sole phosphorylation sites that exhibited a consistently increasing pattern among EGF, PAM and EGF+PAM treatment groups. Interestingly, these two phospho-Tyr sites have been linked to PLCγ signaling ^44^, which, in turn, has been implicated in increased ROS production ^45^. Among the other phosphorylation sites, EGFR(Try1068/1086/1148) were suppressed in response to PAM and PAM+EGF, suggestive of the differential modulation of EGFR phosphorylation sites of the proposed PAM-based treatment modality towards improved anti-cancer efficacy. EGFR(Try1068/1086) and EGFR(Try1148) each mediates the GRB2/STAT3 and GRB2/ERK signaling, respectively, and is oncogenic, with GRB2 being a known therapeutic target of many solid tumors including breast cancers^46^. Thus, by simultaneously suppressing GRB2-mediated pathways and elevating PLCγ signaling, PAM, alone or coupled with EGF, is capable of achieving the observed selectivity against TNBCs via a unique molecular mechanism that cannot be obtained by conventional targeted therapies.

Besides transducing signals, ROS may oxidize DNA that overwhelms the DNA repair apparatus and triggers programmed cell death including apoptosis once entering the cell nucleus ^47^. Thus, ROS accumulation in the nucleus as triggered by EGF in combination with PAM (**Figure 4d**) may contribute to the increased PAM sensitivity of TNBC cells (**Figure 1d-e**). Although some residual signaling and proliferative responses have been noted with EGFR mutations such as V741G ^25,48^, they were not sufficient to support PAM effects on cell viability (**Figure 1j**), further differentiating proliferative signaling responses mediated by EGFR(V741G) from that involving PLCγ, in relation to PAM induced loss of cell viability.

A key feature of our study was the observation that EGFR internalisation and lysosomal degradation was blocked by PAM treatment, allowing EGFR to continue sustained cytoplasm signaling until PAM was removed. This may, at least partially, be attributed to a series of post-translational modification (PTM) events that involve, e.g., orchestrated efforts of phosphorylation and ubiquitination. K48 and/or K63 ubiquitination are required for EGFR endocytosis trafficking and subsequent lysosomal degradation ^49,50^, and reduced EGFR ubiquitination would lead to increased EGFR retention on the plasma membrane. Consistent with this, we observed EGFR accumulation on the cell surface after CAP/PAM treatment using EGF-Alexa 488 pulse-chase uptake assays (**Figure 1i**). This is the standard for monitoring EGFR endocytosis as EGFR antibodies alter receptor trafficking and steady state staining does not show alterations in the kinetics of endocytosis ^24,49–51^.

Among the three TNBC lines examined, EGF showed the optimal synergy with PAM followed by TGFα (**Figure S5g**), the latter of which was reported to elevate EGFR expression at both the transcriptional and translational levels in prostate cancer cells^52^. Thus, we focused on EGFR-relayed signaling in this study to investigate the anti-cancer efficacy of PAM against TNBC cells. However, we do not exclude the possibility of other tyrosine kinase receptors participating in PAM-triggered cell responses, and indeed we saw a PAM response in the BaF/3 hematopoietic cell line that lacks endogenous EGFR and most other known RTKs except for IL2 receptor. In particular, knocking out TGFβR, another cell RTK that plays critical roles in cancer initiation and progression ^53,54^, enhanced the efficacy of PAM in reducing the viability of cancer cells (**Figure 1k**). Besides, SUM-159PT cells exhibit higher expression of FGFR1/2 than other breast cancer cell lines ^55,56^ that can enhance PLCγ phosphorylation ^57,58^; consistent with this, SUM-159PT cells were more sensitive to PAM+FGFs than other cell lines **(Figure S5g)**, suggestive of the involvement of FGFR in PLCγ-relayed signaling in response to PAM treatment. Thus, PAM-triggered cell apoptosis, though being an orchestrated effort from a panel of cell receptors, take actions via PLCγ signaling. In addition, EGFR is selectively expressed in the TNBC cell lines (**Figure 5c**) ^59^, and the sensitivity of cancer cells to PAM treatment is quantitatively related to EGFR activity (**Figure 1k, 5e, 5f**). Thus, EGFR may represent a potential target for this clinical subtype, and an avenue through which the efficacy of plasma therapy could be enhanced.

It is also important to address that PAM-mediated anti-cancer efficacy is dose-dependent ^60^. With the increase of PAM doses, cells can initiate the survival program such as autophagy^61^ and cell cycle arrest^62^, and programmed cell death such as apoptosis^12^, ferroptosis^63,64^ and immunogenic cell death (ICD)^65^. By synergizing with EGF under the examined PAM doses, the apoptotic program is particularly initiated by modulating the differential EGFR phosphorylation sites as we reported in this study.

CAP is a cocktail of reactive species, among which we identified hydroxyl radical (OH), superoxide anion (O_2_^−^), nitric oxide (NO), and (H_2_O_2_) as the leading RONS that mediate the action of PAM as well as its synergy with EGF in reducing the viability of TNBC cells. As CAP generated short-lived reactive species ^66^ and the EGFR monoclonal antibody cetuximab ^67^ could both induce ICD in cancer cells, it is likely that short-lived reactive species from CAP (such as·OH, O_2_^−^, NO) and/or generated from interactions among CAP components (such as O_2_^−^) activated PLCγ signaling via stimulating EGFR(Tyr992/1173). Thus, any individual component of PAM such as O_3_ and H_2_O_2_ is unlikely to function as effectively as the full PAM as an oncotherapy, as also reported by others ^66,68^. Further, the important roles of short-lived reactive species such as OH (half-life 10^−9^s *in vitro* and *in vivo*), O_2_^−^ (half-life 10^−6^s *in vivo*, 10^−3^s *in vitro*), and NO (~1s *in vivo*, seconds to hours *in vitro*) ^69^ in delivering the anti-cancer efficacy of PAM render it difficult to reconstitute PAM that is typically generated via appropriate Ar/He-based plasma source. It is worth mentioning that the lifetime of these reactive species varies with different conditions of the treated solution (i.e., pH, temperature and chemicals inside). However, such variations do not change the classification of these species (i.e., short- or long-lived) and thus do not affect the clinical efficacy and usage of PAM. Indeed, sodium lactate Ringer’s solution has been recommended for generating PAM if applied clinically ^70^, as well as drug release system^71^. The N_2_ and N-containing species observed from OES (**Figure 1b**) originated from the surrounding air, since no further protecting gas and setup for isolating air were employed in our study. The OH formed at the plasma-liquid interface (which has been widely reported in the literature) was mainly generated from the reactions between water molecules and energetic Ar* or electrons. A short and mild plasma exposure or low dose plasma treatment (such as the use of a cold plasma jet for several minutes) may not strongly affect the biomacromolecules (e.g., proteins) in the liquid, while the small molecular compounds (amino acid or even glucose) may consume a small amount of aqueous species. Overall, the plasma-based medium activation strategy does not affect the clinical efficacy and usage of PAM, which has been confirmed in some published studies ^72,73^.

Fabrication of plasma sources feasible for administrating CAP to different types of tumors and tumors at different loci to enable direct contact of the cocktail formula of CAP to the pathological sites is required for the clinical translation of plasma for cancer treatment. In addition, PAM might be made into capsules where the activities of short-lived species such as OH and O_2_^−^ could be maintained by appropriate nanoparticles and thus be used for oncotherapy. In this research, the direct intratumoural injection of PAM had been conducted.

The recent observation that PLCγ could also physically bind to PD-L1 even in EGFR p-Tyr992/1173 mutated cells, resulting in tumor cell proliferation ^74^, suggests that combined EGF and PAM treatment could competitively occupy PLCγ to generate ROS for cytotoxicity, rather than allowing it to drive cell proliferation signaling. This would provide a theoretical basis for the role of plasma in immunotherapy in the future.

We demonstrated in this study that TNBC cells, which have higher levels of EGFR than the corresponding normal tissue (>70% of patient samples) ^75,76^, show enhanced cytotoxic responses to PAM treatment. EGF dramatically enhances the killing of the TNBC cells by PAM, mediated via the EGFR (p-Tyr992/1173)/PLCγ axis that triggers cell apoptosis due to elevated cellular ROS levels. This study opens a new horizon in the clinical use of EGFR ligands, and/or modified forms thereof, as onco-therapeutic agents in conjunction with PAM, which is contradictory to conventional onco-therapeutic strategies relying on EGFR blockage. We identified EGF as a novel ‘drug’ towards enhanced therapeutic efficacy of PAM in cancer treatment, which warrants careful reconsideration of the clinical use of traditional FDA-approved drugs that block EGFR signaling if PAM or other drugs modulating cellular ROS level (e.g., radiation treatment) are also being used for cancer control. Although this study focused on EGFR due to its strong association with TNBC, we also provide preliminary evidence for similar synergism with other RTKs when present in the cells, paving the way for a personalized approach based on tumor-specific RTK profiles, and /or generic downstream signaling targets.

## MATERIALS AND METHODS

### Cell Culture

Cells were cultured at 37 °C incubator with 5% CO_2_ in DMEM with 10% FCS (HEK-293T, SUM-159PT, MCF-7 and MDA-MB-468) or RPMI with 10% FCS (HCC70, MDA-MB-231, T-47D). MCF-10A was cultured in DMEM supplemented with 20 ng/mL EGF, 100 ng/mL Cholera Toxin, 5% horse serum, 10μg/mL insulin, 1% streptomycin and 0.5 mg/mL hydrocortisone. Cells were passaged with trypsin and stored in liquid nitrogen. SUM-159PT cells were purchased from BeNa Culture Collection (Cat# 359533). Other cells were obtained from ATCC (Manassas, USA). Cells were subjected to STR profile validation and routinely checked for mycoplasma.

### Antibodies and reagents

Primary antibodies: EGFR (Cat# sc-120, Santa Cruz, USA), Phospho-EGFR (Tyr992) (Cat# PA5-85794, Invitrogen, USA), T-PLCG1 (Cat# PA5-83347, Invitrogen, USA), Phospho-PLCG1(Tyr783) (Cat# 700044, Thermo-Fisher Scientific, USA), T-AKT (Cat# 4691, CST, USA), Phospho-AKT (Ser473) (Cat# 4060, CST, USA), p44/42 MAPK (Erk1/2) (Cat# 9102, CST, USA), Phospho-p44/42 MAPK (Erk1/2, Thr202/Tyr204) (Cat# 9106, CST, USA), β-actin (Cat# sc-8432, Santa Cruz, USA), D9D5 K48-linkage polyubiquitin (Cat# 8081, CST, USA), GADPH (Cat# AC001, ABclonal, China).

Secondary antibodies: Goat anti-mouse IgG-HPR (Cat# sc-2031, Santa Cruz, USA), goat anti-rabbit IgG H&L-HRP (Cat# ab6721, Abcam, USA). Phalloidin Alexa Fluor 647 for F-actin staining (Cat# A22287, ThermoFisher, USA).

### Reagents

A6355 (ERK1/2 inhibitor, Cat# MFCD09038681, Sigma-Aldrich, USA), A6730 (AKT1/2 inhibitor, Cat# MFCD08705407, Sigma-Aldrich, USA), U73122 (PLCγ inhibitor, Cat# 112648-68-7, Sigma-Aldrich, USA), erlotinib hydrochloride (EGFR tyrosine kinase inhibitor, Cat# ab141930, Abcam, USA), gefitinib (EGFR tyrosine kinase inhibitor, Cat# ab142052, Abcam, USA), cetuximab (EGFR monoclonal antibody, Cmab, Merck Serono, Germany), panitumumab (EGFR monoclonal antibody, Pmab, Amgen, USA), Interleukin-3 (IL-3, Cat# SRP4135-10UG, Sigma, USA), Alexa Fluor 488 EGF complex (Cat# E13345, Thermo-Fisher Scientific, USA), CellROX^®^ (Cat# C10443, Thermo-Fisher Scientific, USA), Hoechst 33342 (Cat# H3570, Thermo-Fisher Scientific, USA), human Epidermal Growth Factor (hEGF, Cat# 62253-63-8, Sigma-Aldrich, USA), human Transforming Growth Factor-α (hTGF-α, Cat# 105186-99-0, Sigma-Aldrich, USA), human Transforming Growth Factor-β1 (hTGF-β1, Cat# T7039, Sigma-Aldrich, USA), Fibroblast Growth Factor 1 (FGF1, Cat# SRP2091, Sigma-Aldrich, USA), Fibroblast Growth Factor 2 (FGF2, Cat# F5392, Sigma-Aldrich, USA), Fibroblast Growth Factor 10 (FGF10, Cat# F8924, Sigma-Aldrich, USA).

### Live-Dead Assay

Cells were plated in a 96-well plate for 24h at 5,000 cells per well. Following incubation of cells with the indicated treatments, propidium iodide (10 μg/mL) and Hoechst 33342 (10 μg/mL) were added 30 min before imaging. Cells were imaged on an IN-Cell Analyzer 2200 (GE Healthcare, Location; 10×objective). Live/dead cell analysis was performed using IN Cell analysis software.

### PAM generation

The set-up of the CAP device and PAM generation are shown in **Figure 1a**. The CAP device was a typical high-frequency (1.7 MHz, 2-6 kV), atmospheric pressure Argon plasma jet (model kINPen 09) developed at the Leibniz-Institute for Plasma Science and Technology (INP Greifswald), Germany. As shown in **Figure 1a**, CAP was generated between central electrode and ring electrode and flowed out of the quartz tube via the Argon gas with a flow rate of 5.0 Standard Liter per Minute (power: < 3.5 W in the hand-held unit). Aliquots of 1.5 mL DMEM (Cat# 11965092, Lot#, 1965453, ThermoFisher Scientific) without serum in 2 mL centrifuge tubes were activated by CAP for 10 min and named ‘10PAM’ here. PAM was then diluted to different concentrations, designated by the percent remaining (e.g., 70% 10PAM refers to 70% concentration at use). After treatment, different concentrations of PAM were used in cultured cells, which were seeded in a 96-well plate.

### RONS measurement and scavengers

A spectrometer (Andor Shamrock SR-500i-A-R) was used to capture the optical emission spectra (OES) of the plasma discharges, reactive species in the gas phase and to validate the formation of secondary chemical species in the treated medium (**Figure 1b**). OES of the CAP was dominated by hydroxyl band (A^2^Σ to X^2^ Π, 0-0, 1-1, 2-2), second positive system emission of N_2_ (C^3^Πµ to B^3^Πg), argon bands and oxygen emission lines ^77^. Chemical scavengers for ROS and RNS, D-mannitol (Sigma-Aldrich, 200 mM), tiron (Chem-Supply Pty Ltd, 20 mM), sodium pyruvate (Sigma-Aldrich, 10 mM), monopotassium (Sigma-Aldrich, 1 mM), uric acid (Sigma-Aldrich, 100 µM) and hemoglobin (Sigma-Aldrich, 20 µM), were employed to trap ·OH, ·O_2_^−^, H_2_O_2_, e^−^, O_3_ and ·NO respectively.

The titanium sulfate method was used to quantify H_2_O_2,_ where a yellow-colored complex was formed with a UV–Vis absorbance at 410 nm following the reaction Ti^4+^ + H_2_O_2_ + 2H_2_O → H_2_TiO_4_ + 4H^+^) ^78^. A multiparameter photometer (Hanna Instruments, HI83399) and a certified O_3_ reagent kit purchased from the supplier were used for O_3_ quantification via the colorimetric method. The Griess Reagent method was used to quantify NO_2_^−^, and a nitrate specific ion electrode was used to quantify NO_3_^−^ where 2 mol/L Ionic Strength Adjuster (NH_4_)_2_SO_4_ was added before detection ^79^.

### Annexin V/PI analysis

HEK-293T, SUM159 or T47D cells were trypsinized into 1.5 mL centrifuge tubes that were cleaned three times using pending buffer. Cells were pelleted and stained with the recommended concentration of Promega Annexin V-FITC Apoptosis Detection Kit (United Bioresearch; Dural NSW, Australia). After staining, cells were re-suspended with pending buffer to 2×10^5^–5×10^5^ cells in 400 μL buffer. 488-Annexin antibody was diluted in 1:40, and cells were incubated for 15 min in darkness. After washing and re-suspending cells 3 times, cells were stained by propidium iodide (PI) at 1:20 dilution. Fluorescence of cells was measured using the Gallios flow cytometer system and analyzed using the Kaluza software (Beckman, Lane Cove NSW, Australia).

### HoloMonitor imager measurements

SUM159 and MDA-MB-231 cells were plated in 96-well plates at 5,000 cells per well for 24h before real-time monitoring. When the cell density was around 30%, the digital holograms of cells were set up using the HoloMonitor M4 Digital Holography Cytometer (Phase Holographic Imaging PHI AB, Lund, Sweden). The results were calculated using Hstudio M4 software (Phase Holographic Imaging PHI AB, Lund, Sweden)

### DeltaVision Ultra High-Resolution Microscope

MDA-MB-468 cells were plated at 5 × 10^5^ per well in an 8-well chamber slide for 24 h with or without 30 min PAM pre-treatment, and EGF-Alexa 488 added to the medium simultaneously. The 488 nm laser of DeltaVision OMX V3 was used to provide the illumination, and pictures were captured on two cameras: Photometrics Cascade (Photometrics, Tucson, USA) and back-illuminated EMCCD cameras (90% QE). Data capture used an Olympus UPlanSApo 10061.4 NA oil objective and standard excitation and emission filter sets (in nm, 488 EX/500–550 EM and 592.5 EX/608–648 EM). 3D-SIM images were sectioned using a 125 nm z-step.

### Metabolic energy ATP measurements

The rates of ATP generation from glycolysis and mitochondria were measured via the Agilent Seahorse XFe96 Analyzer (Agilent Technologies, Santa Clara, CA) as described ^80^. The oxygen consumption rate (OCR) value is used to calculate ATP produced through OXPHOS from mitochondria and the extracellular acidification rate (ECAR) value to calculate ATP generated from glycolysis process. Briefly, 2 × 10^4^ cells of each cell line were seeded in 96-well-plate Seahorse XF cell culture plate at 24 h before the treatment. After the half hour specific pre-treatment, the cells were washed 3 times and cultured in phenol red free DMEM medium (supplemented with 10 mM glucose, 1 mM sodium pyruvate and 2 mM L-glutamine without serum) 1 h at 37 °C in atmospheric CO_2_ incubator. To determine the OCR and ECAR values, the ATP rate assay, including 1.5 μM Oligomycin and 1μM each of Antimycin A and Rotenone, were added during the test process consecutively. ATP production rate was calculated using Wave software. These values were normalized for the cell numbers of each sample, counted by InCell HS6500 following a 10-min incubation with 10 μg ml-1 Hoechst 33342.

### ELISA

The EGFR resting expression level of each cell line were measured by enzyme-linked immunosorbent assay (ELISA), STAR EGFR ELISA Kit (Millipore, Billerica, MA) and followed by manufacturer’s recommended protocol. The cells were plated in 6-well-plates at 5 × 10^5^ per well for 24 h. The adherent cells were washed 3 times with PBS and incubated with 100 μL cold RIPA buffer containing 1% SDS and protease inhibitors for 30 min on ice. The fresh protein lysates were collected by scraping from the plate into 1.5 mL Eppendorf tube and briefly sonicated for 10 seconds. Centrifugation at 12,000 rpm for 10 minutes at 4 °C. The total proteins were in the supernatant and normalized to 1mg/mL based on the Micro BCA assay according to manufacturer’s recommendations (Thermo Fisher Scientific Inc, Rockford, IL). 25 μL of each sample was diluted in the Diluent Buffer to 100 μL and added in the EGFR capture plate. Remaining protein lysates were kept at −80 °C. The standard EGFR protein was added and plates incubated for 2 h at room temperature with shaking, before removing samples and washing with provided washing buffer 4 × 5 min. The plate was then incubated with EGFR detection antibody for 1h, anti-Rabbit IgG HRP conjugate for 45 min, TMB Solution for 30 min in the dark, and Stop Solution, successively, with 4 × 5 min washing between each step. The data was immediately collected by Multiskan™ FC Microplate Photometer (Thermo Fisher Scientific, USA) at 450 nm. Based on the EGFR standard curve, the EGFR level of each sample was calculated.

### Human EGFR Phosphorylation Antibody Array

Human EGFR Phosphorylation Array Kit was purchased (CatLog number ab134005, Abcam, USA). The 2 × 10^7^ cells were seeded in T25 flask for 24 h and were rinsed with PBS and solubilized with 1mL 1× lysis buffer, containing Protease Inhibitor Cocktail and Phosphatase Inhibitor Cocktail Set I and Set II provided in the kit. Lysates were incubated gently on the ice for 30min after transfer to the microfuge tubes, and centrifuged at 14000 × g for 10 min at 4 °C. According to the Micro BCA assay (Thermo Fisher Scientific Inc, Rockford, IL), every sample was normalized to 4 mg/mL and 200 μL diluted in 5% BSA in TBST to 1 mL. The remaining protein lysates were stored at −80 °C fridge. The membranes were placed in the 8-well tray and incubated with 2 ml blocking buffer at room temperature for 1h with shaking. After the blocking buffer removal, 1 ml cell lysate of each sample was placed on the membrane and incubated at room temperature for 2 h. The membranes were washed 3 × 3 min with 2 mL wash buffer I and removed to a container containing 20mL wash buffer II 3 × 5 min. The membranes were incubated with 1mL Biotin-Conjugated anti-EGFR for 2 h at room temperature overnight on an orbital shaker, and washed again using the same previous method. 1.5 ml of HRP-conjugated streptavidin solution was added onto the membrane and incubated at room temperature for 2 h before the same wash steps. Then, 500 μL Detection Buffer comprising 250 μL Detection Buffer C and 250 μL Detection Buffer D was added onto each membrane and the membranes placed on a plastic sheet. The ChemiDoc™ Touch Imaging system (Bio-Rad, CatLog number 5119100, Hercules, CA, USA) was used to take detailed pictures of the array membranes, which were quantified by the Image Lab Software 6.0.1 (Life Science Research, Bio-Rad, USA). The membranes were stored at −20 °C for future reference.

The normalization of each signal followed the protocol recommended by the manufacturer, which contains two steps. One is normalized by the positive controls between two membranes, and the other is to normalize each phosphorylation spot.

Pos (1) = average signal intensity of positive controls on the reference array

Pos (2) = average signal intensity of positive controls on Array 2

X (2) = signal intensity for a particular spot on Array 2

X (N2) = the normalized value for that particular spot on Array 2 X (N2) = X(2) × Pos (1) / Pos (2)

After positive control normalization, phospho-EGFR signals were normalized to the pan EGFR signals that could be calculated according to the following example:

EGFR (1) = average signal intensity of pan EGFR on the reference array

EGFR(2) = average signal intensity of pan EGFR in Array 2

Y845 (2) = average signal intensity of EGFR (Tyr845) in Array 2

Y845 (N2) = normalized signal of EGFR (Tyr845) in Array 2

Y845 (N2) = Y845 (2) × EGFR (1) / EGFR (2)

Therefore, the normalized signal value of each array is calculated. In order to facilitate comparison, we showed the Log2 value. Each group was scanned three times and the SD value was calculated.

### Immunofluorescence microscopy

SUM-159PT, MCF-10A, T47D and MDA-MB-468 cells were plated at 5,000 cells per well for 24h, with or without different PAM treatments. After the treatment for the designated durations, cells were fixed with 4% paraformaldehyde in PBS for 10 min at 4 °C. Cells were first permeabilized in 0.2% Triton X-100 for 5 min and then blocked (3% BSA in PBS) for 1 h, followed by incubation with primary antibodies as indicated. After incubation, cells were washed three times with PBS before the addition of secondary antibodies as indicated, and these were incubated for 1 h at the room temperature. Nuclear DNA was stained with Hoechst 33342 for 5 min at 10 μg/mL. A Delta Vision Elite Live Imaging Microscope (Applied Precision, GE Healthcare Lifesciences, Parramatta NSW, Australia) and softWoRx analysis software (Applied Precision) were used for cell imaging. EGFR immunofluorescence was assessed using Axio Imager Z2 (Zeiss) microscope. High-throughput analysis was performed using an IN Cell 2200 Analyzer and IN Cell analysis software (GE Healthcare).

### Reactive oxygen species (ROS) Assay

Cells were treated with or without PAM for the specified time duration and the CellROX® Reagent was added to cells at a final concentration of 5 μM and incubated for 30 minutes at 37 °C. Medium was removed and cells were washed 3 times with PBS, after which cells were stained with 10 μg/ml Hoechst 33342, a nuclear counterstain.

### CRISPR/Cas9 Gene Editing

Custom sgRNAs for human EGFR, FGFR and TGFβR gene were designed using the MIT CRISPR Design website (https://www.benchling.com/crispr/) with the sequence of EGFR (NM_005228.5), FGFR (NM_023110.3), TGFβR (NM_001024847.2). This website predicts both on-target sequences and off-target possibilities. We selected the best scoring guide sequences in the EGFR protein-coding region as sgRNA_1 (GAATTCGCTCCACTGTGTTG) and sgRNA_2 (CGATCTCCACATCCTGCCGG); FGFR protein-coding region as sgRNA_1 (GTTGCCCGCCAACAAAACAG) and sgRNA_2 (GCGGTGGCTGAAAAACGGGA); TGFβR protein-coding region as sgRNA (TGGGCAGTCCTATTACAGCT). A scramble (SC) sgRNA (CCATATCGGGGCGAGACATG) was used as a parental control. Guide oligonucleotides were phosphorylated, annealed, and cloned into the BsmBI site of the lentiCRISPR v2 vector (Addgene, 52961, kindly provided by Feng Zhang), according to the Zhang laboratory protocol (F. Zhang lab, MIT, Cambridge, MA, USA). All plasmid constructs were verified by sequencing.

### Plasmid Construction, Lentiviral Production and transduction

Lentiviral particles were produced by transient transfection of Phoenix-ECO cells (CRL-3214) using TransIT®-LT1 Reagent (Mirus Bio LLC, Madison, WI, USA). The lentiCRISPR construct or the pLJM1-EGFP plasmid (Addgene plasmid #19319, a gift from David Sabatini) was co-transfected with pMD2.G (Addgene plasmid #12259) and psPAX2 (Addgene plasmid #12260, both kindly provided by Didier Trono, EPFL, Lausanne, Switzerland). Lentiviral particles were collected at 36 and 72 h and then purified with 10% Lenti-X Concentrator® (Clontech, Mountain View, CA, USA). The virus copy number was determined by Q-PCR. A total of 1 × 10^6^ cells was plated in a 6-cm dish and treated with EGFR targeting lentivirus at an MOI = 5 for 48 hours. The culture medium was replaced and antibiotic selected with 2.5 ug/mL puromycin for 48hours. The cell line transfected with sgRNA_1 virus refers to KO1 in this study, whereas sgRNA_2 virus transfected cell line refers to KO2. The protocol of gene recombinant and infectious biological materials experiment was approved by the Institutional Biosafety Committee (G-109-059 and G-108-024) at Taipei Medical University, Taipei, Taiwan.

### Animal Trials

*In vivo* analysis was undertaken using intratumoural injection of PAM generated by a home-made plasma source^81^. The experiment was performed twice. PAM was generated by setting the distance between the CAP nozzle and the medium surface to 13 mm, the peak-to-peak electrode voltage to 1.1 kV, the sinusoidal wave frequency to 8.8 kHz, the Helium gas flow rate to 1 L/min, and exposing the medium to CAP for 5 min.

SUM159PT cells (1 × 10^6^) suspended in phosphate buffer saline (PBS) were injected subcutaneously in the right forelimbs of 50 female BALB/c mice aged 4 weeks with the weights of 16 ± 2 g on the first day. Nine mice carrying SUM159PT tumors which had grown to a similar size 3 weeks later were recruited into each study. The mice were evenly divided into 3 groups, i.e., SUM159PT_control group (without PAM treatment), SUM159PT_PAM group (receiving PAM), and SUM159PT_PAM+Afa group (receiving CAP and afatinib). The first treatment was performed when tumor sizes reached 5 ± 0.5 mm. Tumor diameters were measured using vernier caliper. Mice were anesthetized with ketamine (concentration is 10 mg/ml) intraperitoneally before each treatment. The injection volume was 10 μL/g of the mouse body weight. PAM was subcutaneously injected at two sites of the tumor for each mouse, with 100 μL/site. This treatment was repeated every 6 days. Tumors were dissected after the sacrifice of the mice on the 19^th^ day. One mouse from the PAM + afatinib (10 μM) group was dead by the time the mice were sacrificed in the first experiment (pilot study).

### Statistical analyses

Most statistical tests of this study were performed by Graph Pad Prism V7. Group-wise analysis was performed using students’ two-tailed t-tests. Mean +/− standard error of the mean (SEM) plotted unless otherwise specified. p value > 0.05 is considered insignificant, “*” represents a p value < 0.05, “**” represents a p value < 0.01, “***” represents a p value < 0.001 and “****” represents a p value < 0.0001.

## Ethical Approval and Consent to participate

The animal experiment complied with the ethical regulations on the use of animal and animal-derived materials and approved by the Animal Laboratory Center of Jiangnan University. Use of human cell lines was approved by the Human Research Ethics Committee of Queensland University of Technology.

## Supporting information

Supplementary Figures Wang et al PAM_EGFR_TNBC

## Acknowledgements

This study was funded by a manuscript grant from the Institute of Health and Biomedical Innovation, Queensland University of Technology, Australia. Funding was also received from the National Natural Science Foundation of China (Grant No. 81972789), Fundamental Research Funds for the Central Universities (Grant No. JUSRP22011), the National Health and Medical Research Council, Australia (APP1187328, APP1165208, APP1187328) and the National Breast Cancer Foundation, Australia (CBC-10-009). These funding sources played no role in the writing of the manuscript or the decision to submit it for publication. The Translational Research Institute receives support from the Australian Federal Government. The authors would like to thank Tony Blick for data analysis, Laura Croft and Jaimie Mulders for Holomonitor data collection and analysis, Sam Beard for high resolution DeltaVision data collection, Katia Bazaka for providing the kINPen 09, Nan Nan for conducting ubiquitination assay, Peiyu Han for helping in the animal experiments.

## Authors’ contributions

Conceptualization: EWT, FS, DR, KO, XD, PW. Methodology: PW, JHG, CHL, AWB, FBF, DR, XD, FS, LZ, LY. Investigation: PW, RWZ, RSZ, XD, JL, AR. Writing-Original Draft: PW, XD. Writing-Review & Editing: all. Funding Acquisition: EWT, XD, KO. Resources: KO, XD, DR, CHL, AWB, FBF, FS. Supervision: EWT, GL, XD, DR, FS, KO

## References

1 Waks, A. G. & Winer, E. P. Breast Cancer Treatment: A Review. JAMA 321, 288–300, doi:10.1001/jama.2018.19323 (2019).

2 Jiang, Y. Z. et al. Genomic and Transcriptomic Landscape of Triple-Negative Breast Cancers: Subtypes and Treatment Strategies. Cancer Cell 35, 428–440.e425, doi:10.1016/j.ccell.2019.02.001 (2019).

3 Keidar, M. et al. Cold atmospheric plasma in cancer therapy. Physics of Plasmas 20, 057101 (2013).

4 Woedtke, T. v., Emmert, S., Metelmann, H.-R., Rupf, S. & Weltmann, K.-D. Perspectives on cold atmospheric plasma (CAP) applications in medicine. Phys Plasmas 27, 070601 (2020).

5 Brany, D., Dvorska, D., Halasova, E. & Skovierova, H. Cold Atmospheric Plasma: A Powerful Tool for Modern Medicine. Int J Mol Sci 21, doi:10.3390/ijms21082932 (2020).

6 Malyavko, A. et al. Cold atmospheric plasma cancer treatment, direct versus indirect approaches. Mater Adv, 1494–1505 (2020).

7 Yan, D. et al. Cold Atmospheric Plasma Cancer Treatment, a Critical Review. Appl. Sci 11, 7757 (2021).

8 Tanaka, H. et al. Plasma-Treated Solutions (PTS) in Cancer Therapy. Cancers (Basel) 13, 1737, doi:10.3390/cancers13071737 (2021).

9 Dai, X., Bazaka, K., Richard, D. J., Thompson, E. R. W. & Ostrikov, K. K. The Emerging Role of Gas Plasma in Oncotherapy. Trends Biotechnol 36, 1183–1198, doi:10.1016/j.tibtech.2018.06.010 (2018).

10 Tornin, J., Labay, C., Tampieri, F., Ginebra, M.-P. & Canal, C. Evaluation of the effects of cold atmospheric plasma and plasma-treated liquids in cancer cell cultures. Nature Protocols 16, 2826–2850 (2021).

11 Chen, Z. et al. Micro-sized cold atmospheric plasma source for brain and breast cancer treatment. Plasma Medicine 8, 203–215 (2018).

12 Xiang, L., Xu, X., Zhang, S., Cai, D. & Dai, X. Cold atmospheric plasma conveys selectivity on triple negative breast cancer cells both in vitro and in vivo. Free Radic Biol Med 124, 205–213 (2018).

13 Xu, X. et al. Quantitative assessment of cold atmospheric plasma anti- cancer efficacy in triple-negative breast cancers. Plasma Process Polym 15, 1800052 (2018).

14 Wang, P. et al. Epithelial-to-Mesenchymal Transition Enhances Cancer Cell Sensitivity to Cytotoxic Effects of Cold Atmospheric Plasmas in Breast and Bladder Cancer Systems. Cancers 13, 2889 (2021).

15 Xia, L., et al. EGFR-targeted CAR-T cells are potent and specific in suppressing triple-negative breast cancer both in vitro and in vivo. Clin Transl Immunol 9, e1135 (2020).

16 Yi, Y. W. et al. Inhibition of the PI3K/AKT pathway potentiates cytotoxicity of EGFR kinase inhibitors in triple-negative breast cancer cells. J Cell Mol Med 17, 648–656, doi:10.1111/jcmm.12046 (2013).

17 Yi, Y. W. et al. Dual inhibition of EGFR and MET induces synthetic lethality in triple-negative breast cancer cells through downregulation of ribosomal protein S6. Int J Oncol 47, 122–132, doi:10.3892/ijo.2015.2982 (2015).

18 Xia, L. et al. Targeting Triple-Negative Breast Cancer with Combination Therapy of EGFR CAR T Cells and CDK7 Inhibition. Cancer Immunol Res 9, 707–722 (2021).

19 Ghosh, S. et al. PD-L1 recruits phospholipase C and enhances tumorigenicity of lung tumors harboring mutant forms of EGFR. Cell Rep 35, 109181 (2021).

20 Tan, X., Lambert, P. F., Rapraeger, A. C. & Anderson, R. A. Stress-Induced EGFR Trafficking: Mechanisms, Functions, and Therapeutic Implications. Trends Cell Biol 26, 352–366, doi:10.1016/j.tcb.2015.12.006 (2016).

21 Adhikari, B. et al. Cold plasma seed priming modulates growth, redox homeostasis and stress response by inducing reactive species in tomato (Solanum lycopersicum). Free Radic Biol Med 156, 57–69, doi:10.1016/j.freeradbiomed.2020.06.003 (2020).

22 Shen, L. et al. Metabolic reprogramming in triple-negative breast cancer through Myc suppression of TXNIP. Proc Natl Acad Sci U S A 112, 5425–5430, doi:10.1073/pnas.1501555112 (2015).

23 Athar, A. et al. ArrayExpress update - from bulk to single-cell expression data. Nucleic Acids Res 47, D711–D715, doi:10.1093/nar/gky964 (2019).

24 Chew, H. Y. et al. Endocytosis Inhibition in Humans to Improve Responses to ADCC-Mediating Antibodies. Cell 180, 895–914 e827, doi:10.1016/j.cell.2020.02.019 (2020).

25 Walker, F. et al. Activation of the Ras/mitogen-activated protein kinase pathway by kinase-defective epidermal growth factor receptors results in cell survival but not proliferation. Mol Cell Biol 18, 7192–7204, doi:10.1128/mcb.18.12.7192 (1998).

26 Peter, M. E. Programmed cell death: Apoptosis meets necrosis. Nature 471, 310–312, doi:10.1038/471310a (2011).

27 Dai, X., Wang, D. & Zhang, J. Programmed cell death, redox imbalance, and cancer therapeutics. Apoptosis 26, 385–414, doi:10.1007/s10495-021-01682-0 (2021).

28 Zhou, R. et al. Underwater microplasma bubbles for efficient and simultaneous degradation of mixed dye pollutants. Sci Total Environ 750, 142295 (2021).

29 Salazar-Cavazos, E. et al. Multisite EGFR phosphorylation is regulated by adaptor protein abundances and dimer lifetimes. Mol Biol Cell 31, 695–708, doi:10.1091/mbc.E19-09-0548 (2020).

30 Wang, C. et al. EGFR-mediated tyrosine phosphorylation of STING determines its trafficking route and cellular innate immunity functions. EMBO J 39, e104106, doi:10.15252/embj.2019104106 (2020).

31 Dai, X., Cheng, H., Bai, Z. & Li, J. Breast Cancer Cell Line Classification and Its Relevance with Breast Tumor Subtyping. J Cancer 8, 3131–3141, doi:10.7150/jca.18457 (2017).

32 Pouliot, N., Nice, E. C. & Burgess, A. W. Laminin-10 mediates basal and EGF-stimulated motility of human colon carcinoma cells via alpha(3)beta(1) and alpha(6)beta(4) integrins. Exp Cell Res 266, 1–10, doi:10.1006/excr.2001.5197 (2001).

33 Hanahan, D. Hallmarks of Cancer: New Dimensions. Cancer Discov 12, 31–46, doi:10.1158/2159-8290.CD-21-1059 (2022).

34 Hanahan, D. & Weinberg, R. A. Hallmarks of cancer: the next generation. Cell 144, 646–674, doi:10.1016/j.cell.2011.02.013 (2011).

35 Mao, Y., Ma, J., Xia, Y. & Xie, X. The Overexpression of Epidermal Growth Factor (EGF) in HaCaT Cells Promotes the Proliferation, Migration, Invasion and Transdifferentiation to Epidermal Stem Cell Immunophenotyping of Adipose-Derived Stem Cells (ADSCs). Int J Stem Cells 13, 93–103, doi:10.15283/ijsc18146 (2020).

36 Sic You, K., Yi, Y. W., Cho, J., Park, J.-S. & Seong, Y.-S. Potentiating Therapeutic Effects of Epidermal Growth Factor Receptor Inhibition in Triple-Negative Breast Cancer. Pharmaceuticals (Basel) 14, 589, doi:10.3390/ph14060589 (2021).

37 Cursons, J. et al. Stimulus-dependent differences in signalling regulate epithelial-mesenchymal plasticity and change the effects of drugs in breast cancer cell lines. Cell Communication and Signaling 13, 1–21 (2015).

38 Acevedo-Díaz, A., Morales-Cabán, B. M., Zayas-Santiago, A., Martínez-Montemayor, M. M. & Suárez-Arroyo, I. J. SCAMP3 Regulates EGFR and Promotes Proliferation and Migration of Triple-Negative Breast Cancer Cells through the Modulation of AKT, ERK, and STAT3 Signaling Pathways. Cancers 14, 2807 (2022).

39 Lyu, N. et al. in Exploration. 20210176 (Wiley Online Library).

40 Gunes, S. et al. Platinum nanoparticles inhibit intracellular ROS generation and protect against cold atmospheric plasma-induced cytotoxicity. Nanomedicine 36, 102436, doi:10.1016/j.nano.2021.102436 (2021).

41 Weng, M. S., Chang, J. H., Hung, W. Y., Yang, Y. C. & Chien, M. H. The interplay of reactive oxygen species and the epidermal growth factor receptor in tumor progression and drug resistance. J Exp Clin Cancer Res 37, 61, doi:10.1186/s13046-018-0728-0 (2018).

42 Truong, T. H. & Carroll, K. S. Redox regulation of epidermal growth factor receptor signaling through cysteine oxidation. Biochemistry 51, 9954–9965, doi:10.1021/bi301441e (2012).

43 Bae, Y. S. et al. Epidermal growth factor (EGF)-induced generation of hydrogen peroxide. Role in EGF receptor-mediated tyrosine phosphorylation. J Biol Chem 272, 217–221 (1997).

44 Sturla, L. M. et al. Requirement of Tyr-992 and Tyr-1173 in phosphorylation of the epidermal growth factor receptor by ionizing radiation and modulation by SHP2. J Biol Chem 280, 14597–14604, doi:10.1074/jbc.M413287200 (2005).

45 Zhang, J., et al. ROS and ROS-mediated cellular signaling. Oxid Med Cell Longev 2016, 4350965 (2016).

46 Chen, D. e\ t al. miR-27b-3p inhibits proliferation and potentially reverses multi-chemoresistance by targeting CBLB/GRB2 in breast cancer cells. Cell Death Dis 9, 188, doi:10.1038/s41419-017-0211-4 (2018).

47 Perillo, B. et al. ROS in cancer therapy: the bright side of the moon. Exp Mol Med 52, 192–203, doi:10.1038/s12276-020-0384-2 (2020).

48 Walker, F., Hibbs, M. L., Zhang, H. H., Gonez, L. J. & Burgess, A. W. Biochemical characterization of mutant EGF receptors expressed in the hemopoietic cell line BaF/3. Growth Factors 16, 53–67, doi:10.3109/08977199809017491 (1998).

49 Huang, F. et al. Lysine 63-linked polyubiquitination is required for EGF receptor degradation. Proc Natl Acad Sci U S A 110, 15722–15727, doi:10.1073/pnas.1308014110 (2013).

50 Liu, Z. et al. The ubiquitin-specific protease USP2a prevents endocytosis-mediated EGFR degradation. Oncogene 32, 1660–1669, doi:10.1038/onc.2012.188 (2013).

51 Simpson, F. et al. SH3-domain-containing proteins function at distinct steps in clathrin-coated vesicle formation. Nat Cell Biol 1, 119–124, doi:10.1038/10091 (1999).

52 Seth, D., Shaw, K., Jazayeri, J. & Leedman, P. J. Complex post-transcriptional regulation of EGF-receptor expression by EGF and TGF-alpha in human prostate cancer cells. Br J Cancer 80, 657–669, doi:10.1038/sj.bjc.6690407 (1999).

53 Gomez-Gil, V. Therapeutic Implications of TGFbeta in Cancer Treatment: A Systematic Review. Cancers (Basel) 13, doi:10.3390/cancers13030379 (2021).

54 Liu, M. et al. TGF-beta suppresses type 2 immunity to cancer. Nature, 115–120, doi:10.1038/s41586-020-2836-1 (2020).

55 Turner, N. et al. FGFR1 amplification drives endocrine therapy resistance and is a therapeutic target in breast cancer. Cancer Res 70, 2085–2094 (2010).

56 Sharpe, R. et al. FGFR signaling promotes the growth of triple-negative and basal-like breast cancer cell lines both in vitro and in vivo. Clin Cancer Res 17, 5275–5286 (2011).

57 Cross, M. J. et al. The Shb adaptor protein binds to tyrosine 766 in the FGFR-1 and regulates the Ras/MEK/MAPK pathway via FRS2 phosphorylation in endothelial cells. Mol Biol Cell 13, 2881–2893 (2002).

58 Melnik, B. C., Schmitz, G. & Zouboulis, C. C. Anti-acne agents attenuate FGFR2 signal transduction in acne. J Invest Dermatol 129, 1868–1877 (2009).

59 Dai, X., Xiang, L., Li, T. & Bai, Z. Cancer Hallmarks, Biomarkers and Breast Cancer Molecular Subtypes. J Cancer 7, 1281–1294, doi:10.7150/jca.13141 (2016).

60 Dai, X., Zhang, Z., Zhang, J. & Ostrikov, K. Dosing: the key to precision plasma oncology. Plasma Process Polym 17, e1900178 (2019).

61 Miao, Y. et al. Cold atmospheric plasma increases IBRV titer in MDBK cells by orchestrating the host cell network. Virulence 12, 679–689, doi:10.1080/21505594.2021.1883933 (2021).

62 Hua, D. et al. Cold atmospheric plasma selectively induces G0/G1 cell cycle arrest and apoptosis in AR-independent prostate cancer cells. J Cancer 12, 5977–5986, doi:10.7150/jca.54528 (2021).

63 Furuta, T., Shi, L. & Toyokuni, S. Non-thermal plasma as a simple ferroptosis inducer in cancer cells: A possible role of ferritin. Pathol Int 68, 442–443, doi:10.1111/pin.12665 (2018).

64 Bekeschus, S. et al. xCT (SLC7A11) expression confers intrinsic resistance to physical plasma treatment in tumor cells. Redox Biol 30, 101423, doi:10.1016/j.redox.2019.101423 (2020).

65 Lin, A. et al. Non-Thermal Plasma as a Unique Delivery System of Short-Lived Reactive Oxygen and Nitrogen Species for Immunogenic Cell Death in Melanoma Cells. Adv Sci (Weinh) 6, 1802062, doi:10.1002/advs.201802062 (2019).

66 Lin, A. et al. Non-Thermal Plasma as a Unique Delivery System of Short-Lived Reactive Oxygen and Nitrogen Species for Immunogenic Cell Death in Melanoma Cells. Adv Sci 6, 1802062, doi:10.1002/advs.201802062 (2019).

67 Pozzi, C. et al. The EGFR-specific antibody cetuximab combined with chemotherapy triggers immunogenic cell death. Nat Med 22, 624–631, doi:10.1038/nm.4078 (2016).

68 Zhou, R. W. et al. Cold atmospheric plasma activated water as a prospective disinfectant: the crucial role of peroxynitrite. Green Chem 20, 5276–5284 (2018).

69 Sun, J. et al. A hybrid plasma electrocatalytic process for sustainable ammonia production. Energy Environ Sci 14, 865–872 (2021).

70 Mateu-Sanz, M. et al. Cold Plasma-Treated Ringer’s Saline: A Weapon to Target Osteosarcoma. Cancers (Basel) 12, 227, doi:10.3390/cancers12010227 (2020).

71 Tu, L., et al. in Exploration. 20210023 (Wiley Online Library).

72 Yan, D. et al. Stabilizing the cold plasma-stimulated medium by regulating medium’s composition. Sci Rep 6, 26016, doi:10.1038/srep26016 (2016).

73 Chauvin, J., Judee, F., Yousfi, M., Vicendo, P. & Merbahi, N. Analysis of reactive oxygen and nitrogen species generated in three liquid media by low temperature helium plasma jet. Sci Rep 7, 4562, doi:10.1038/s41598-017-04650-4 (2017).

74 Ghosh, S. et al. PD-L1 recruits phospholipase C and enhances tumorigenicity of lung tumors harboring mutant forms of EGFR. Cell reports 35, 109181 (2021).

75 Jiang, W., Wang, X., Zhang, C., Xue, L. & Yang, L. Expression and clinical significance of MAPK and EGFR in triple-negative breast cancer. Oncol Lett 19, 1842–1848, doi:10.3892/ol.2020.11274 (2020).

76 Huang, T., Xiang, J., Wang, Y. & Tuo, Y. Changes of EGFR and SMC4 expressions in triple-negative breast cancer and their early diagnostic value. Gland Surg 10, 1118–1124, doi:10.21037/gs-21-119 (2021).

77 Reuter, S., von Woedtke, T. & Weltmann, K.-D. The kINPen—a review on physics and chemistry of the atmospheric pressure plasma jet and its applications. J Phys D Appl Phys 51, 233001, doi:10.1088/1361-6463/aab3ad (2018).

78 Zhou, R. et al. Sustainable plasma-catalytic bubbles for hydrogen peroxide synthesis. Green Chem 23, 2977 (2021).

79 Sun, J. et al. A hybrid plasma electrocatalytic process for sustainable ammonia production. Energy & Environmental Science 14, 865–872 (2021).

80 Bhatia, S., Thompson, E. W. & Gunter, J. H. Studying the Metabolism of Epithelial-Mesenchymal Plasticity Using the Seahorse XFe96 Extracellular Flux Analyzer. Methods Mol Biol 2179, 327–340, doi:10.1007/978-1-0716-0779-4_25 (2021).

81 Zhou, R. et al. Sustainable plasma-catalytic bubbles for hydrogen peroxide synthesis. Green Chemistry 23, 2977–2985 (2021).

