## Supplementary Figures Wang et al PAM_EGFR_TNBC for "Epidermal Growth Factor potentiates EGFR(Y992/1173)-mediated therapeutic response of triple negative breast cancer cells to cold atmospheric plasma-activated medium"

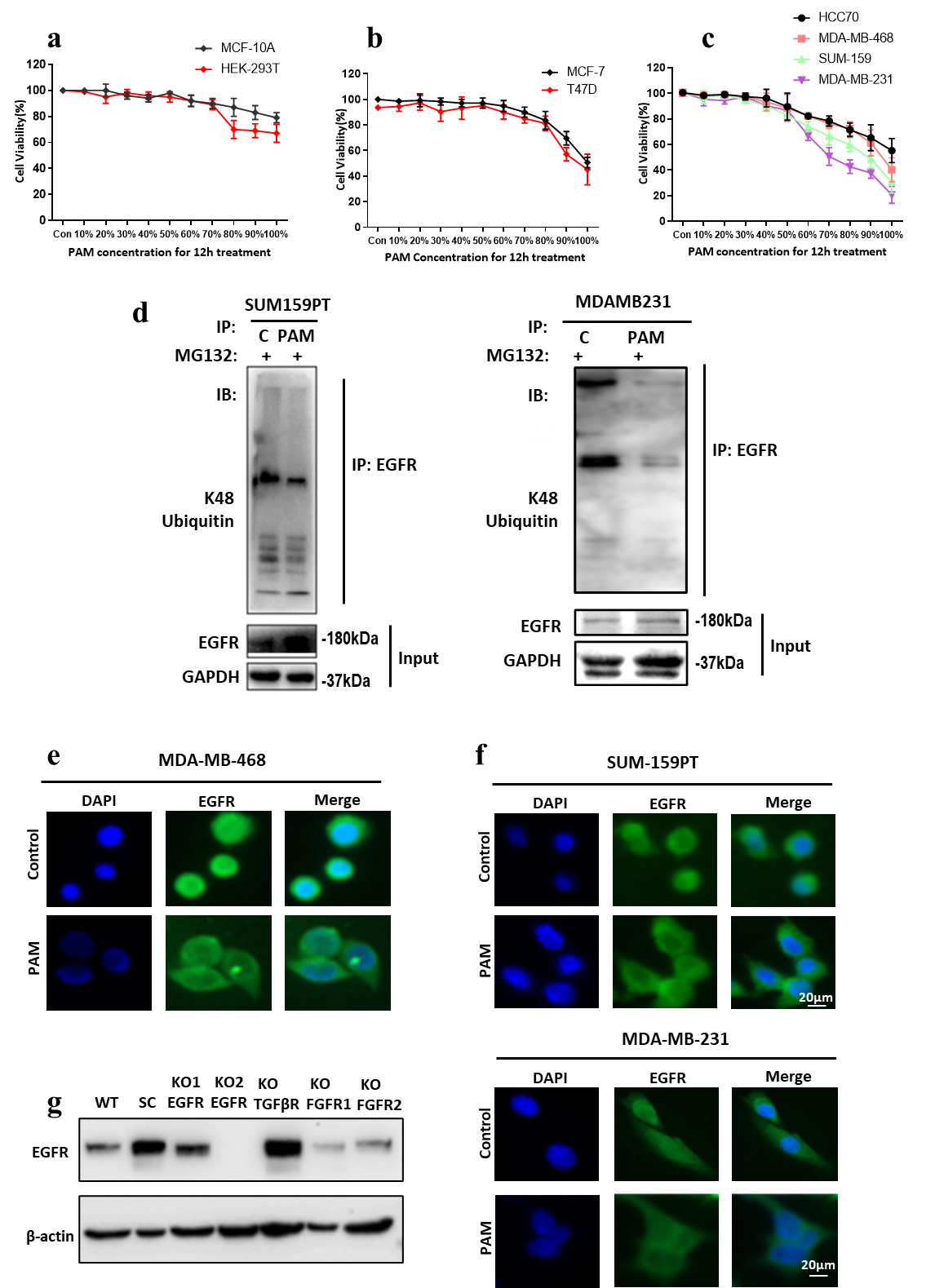

**Supplementary Figure 1. Additional data of Figure 1 (part 1).** Cell viabilities when treated with different concentrations of PAM for 12 h in **a** MCF-10A, HEK-293T, **b** MCF-7, T47D, **c** HCC70, MDA-MB-468, SUM-159 and MDA-MB-231 cells. Cells were plated in 96-well-plate in 5,000 cells per well for 24 h, and treated with 0%, 10%, 20%, 30%, 40%, 50%, 60%, 70%, 80%, 90% and 100% of 10PAM for 12 h. The cells were stained by Hoechst 33342 and PI and cell viability determined. **d** EGFR K48-ubiquitination levels in SUM159-PT and MDA-MB-231 cells with or without PAM treatment. **e** EGFR cellular distribution in MDA-MB-468 cells with or without PAM treatment. **f** EGFR cellular distribution in SUM159-PT and MDA-MB-231 as per **e. g** Knockout efficiency of EGFR, TGFβ, FGFR mutants. Western blot results of wild type MDA-MB-231 (WT), Vector control (SC), EGFR knockout cell lines (KO1 EGFR and KO2 EGFR), TGFβR knockout cell lines (TGFβR KO), FGFR knockout cell lines (KO FGFR1 and KO FGFR2). EGFR antibody (1:1000) was used.

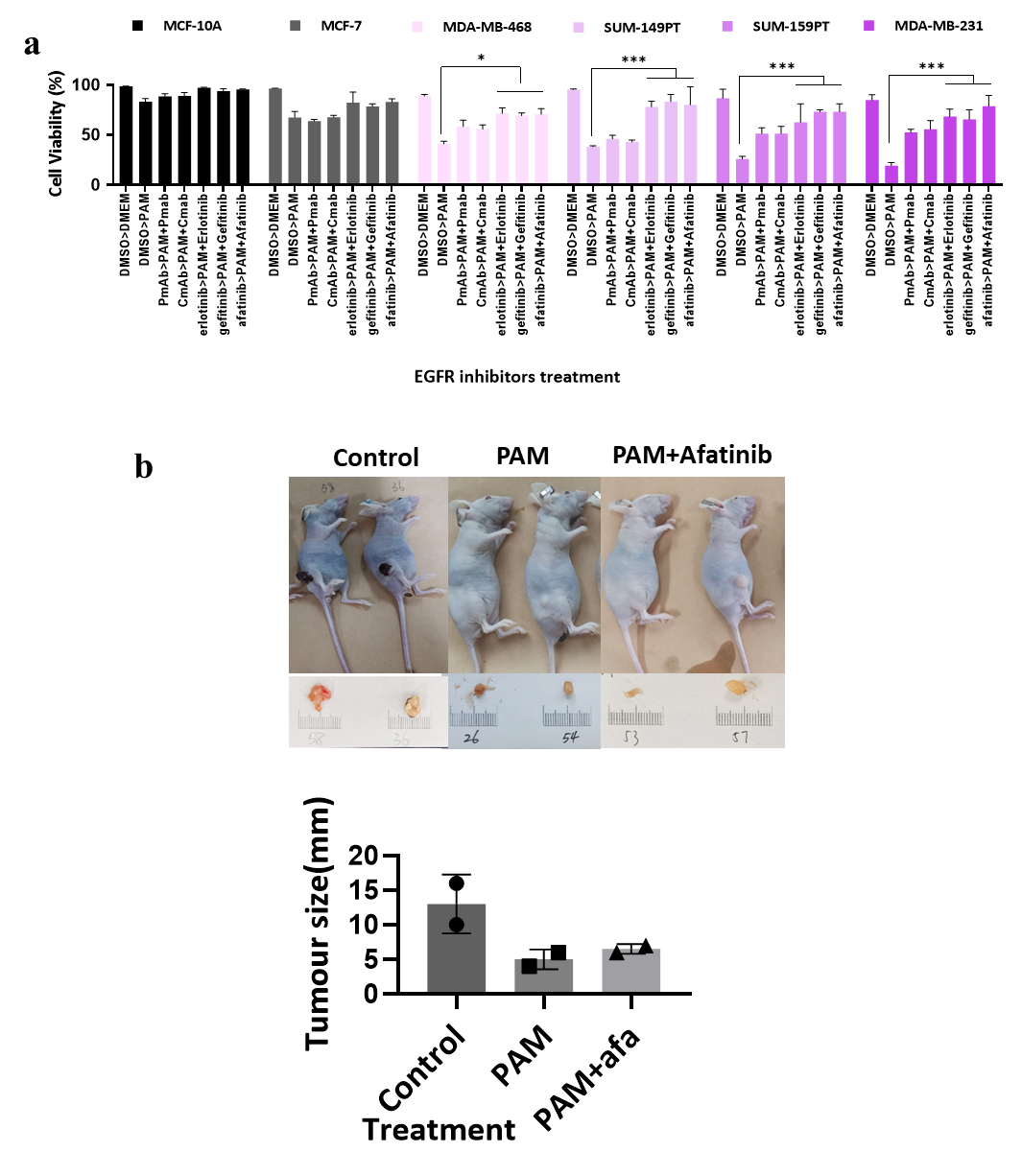

**Supplementary Figure 2. Additional data of Figure 1 (part 2).** **a** Viabilities of MCF-10A, MCF-7, MDA-MB-468, SUM-149PT, SUM-159PT and MDA-MB-231 cells in response to 100% 10PAM treatment and pre-treated with various EGFR inhibitors. Cells were plated in 96-well-plates for 24 h, pre-treated with or without the DMSO (50 ng/mL), PmAb (50 ng/mL), CmAb (50 ng/mL), erlotinib (1 nM), gefitinib (17 nM) or afatinib (10 nM) for 1 h, and then treated with the 100% PAM with these inhibitors for 12 h. N=3. **b** Mice images and tumor sizes from the pilot *in vivo* assay of the control group, CAP treatment group, and CAP plus afatinib treatment group among SUM159-PT tumors.

**Video data is attached separately.**

**Supplementary Figure 3. Additional data of Figure 2 (part 1).** Holomonitor videos of MDA-MB-231 cells in response to EGF, PAM, EGF+PAM treatments. Cells were plated in 96-well-plates at 5,000 cells per well for 24 h, and treated with or without EGF, 50% PAM, EGF+50% PAM for 3 days. During the treatment process, photos of cells were taken every 5min and videos were generated by Holomonitor. The pseudo-color indicates the cell height.

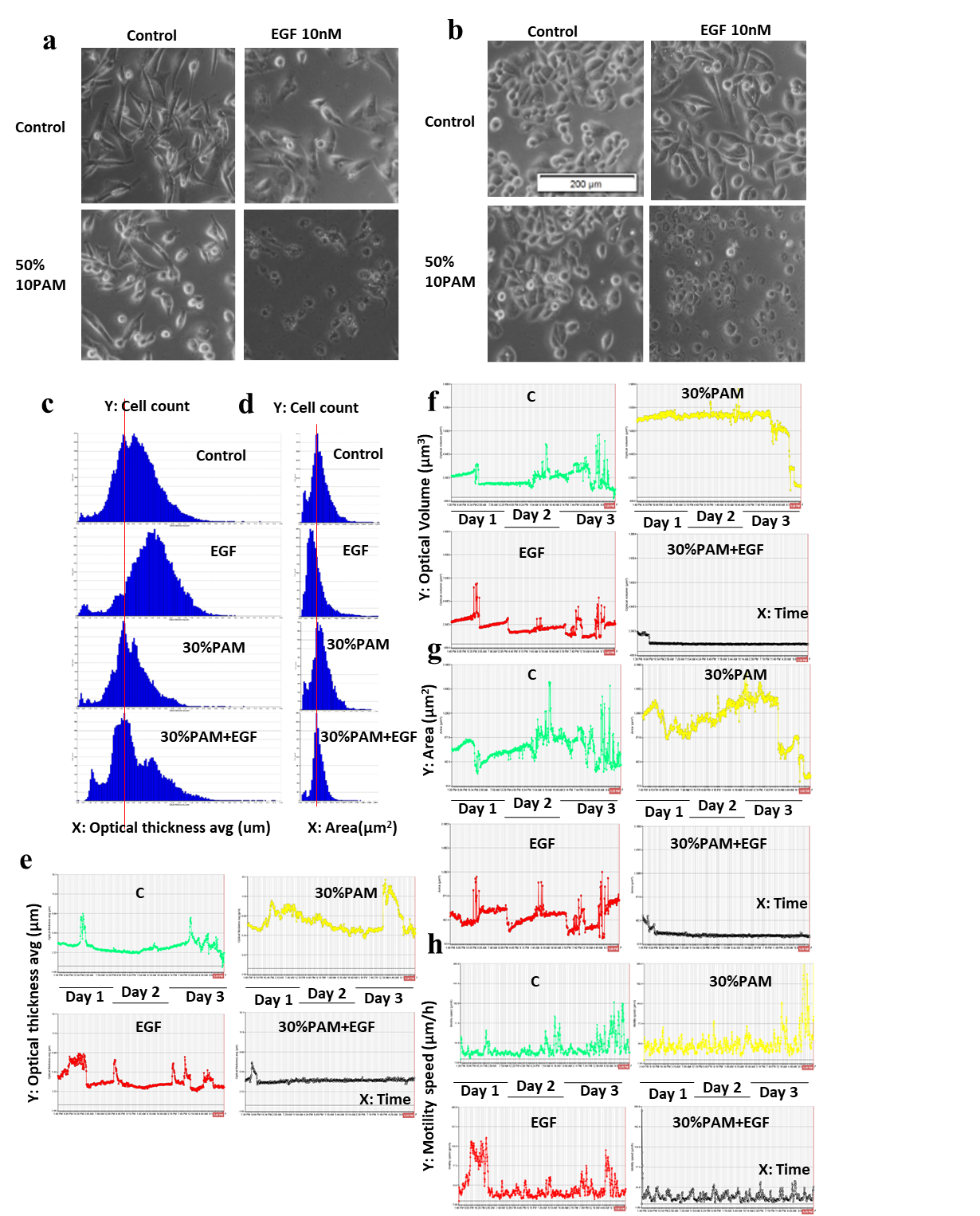

**Supplementary Figure 4. Additional data of Figure 2 (part 2).** Morphology images of (**a**) MDA-MB-231 and (**b**) MDA-MB-468 cells in response to EGF, PAM, EGF+PAM treatments. Cells were seeded in 96-well-plates at 5,000 cells per well for 24 h, and treated with or without EGF, 50% PAM, EGF+50% PAM for 3 days. Photos of cell morphology were taken. Histograms showing relationships (**c**) between average optical thickness (μm) (X axis) and cell count (Y axis), and (**d**) between area (μm^2^) (X axis) and cell count (Y axis) of MDA-MB-468 cells in response to EGF, PAM, EGF+PAM treatments. Curve charts showing the (**e**) optical thickness, (**f**) optical volume (μm^3^), (**g**) area (μm^2^), and (**h**) motility speed (μm/h) changes over 3 days of MDA-MB-468 cells in response to EGF, PAM, EGF+PAM treatments. 10 cells were randomly chosen for the cell morphology analysis. The dynamics changes of the average of 10 cells were shown.

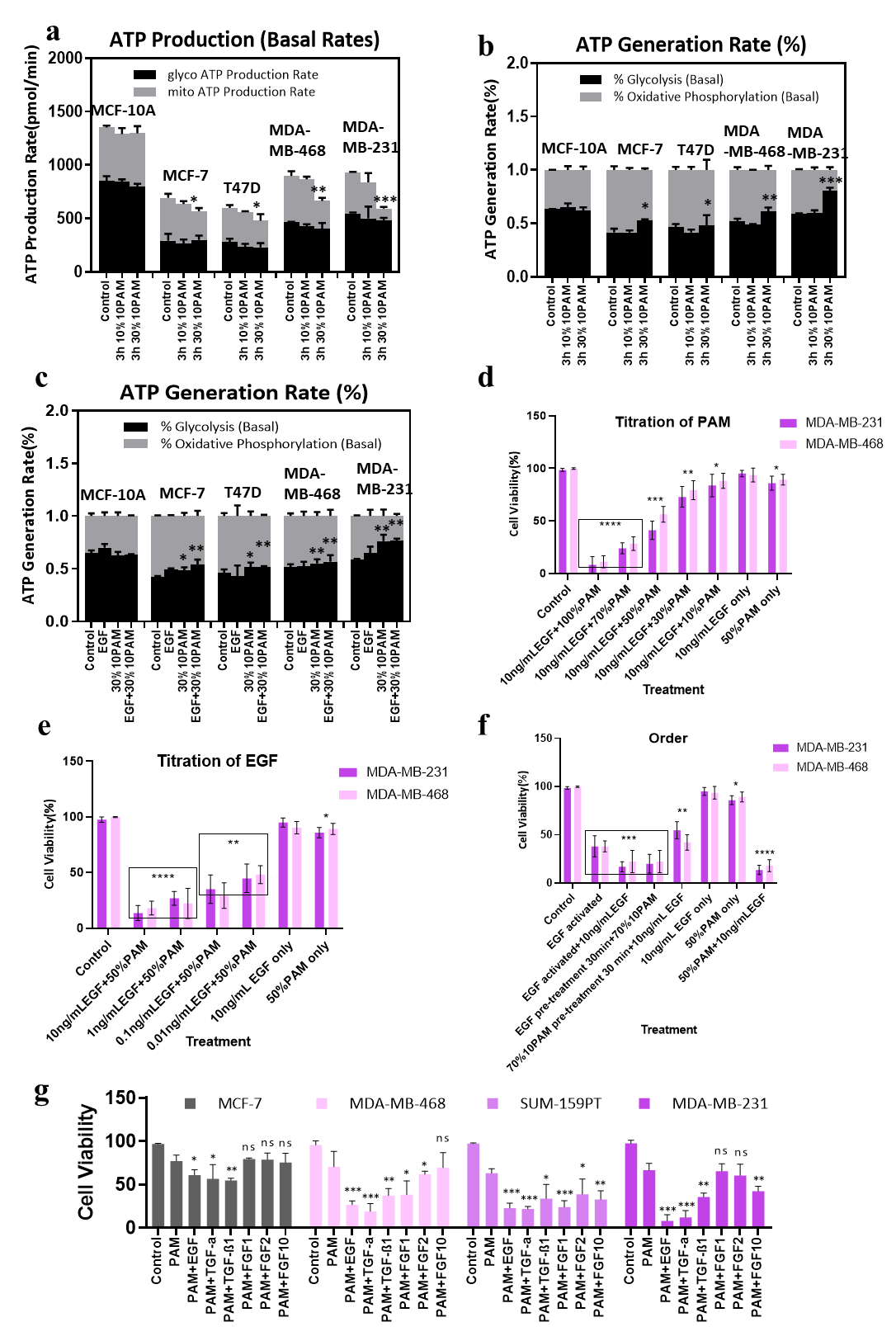

**Supplementary Figure 5. Additional data of Figure 2 (part 3). a** ATP production rates of different cell lines in response to PAM of different PAM concentrations. Percentage of ATP production rates of different cell lines in response to (**b**) PAM of different concentrations, and (**c**) EGF, PAM, EGF+PAM treatments. MCF-10A, MCF-7, T47D, MDA-MB-468 and MDA-MB-231 cells were seeded at 2 × 10^4^ cells per well in 96-well-plate Seahorse XF cell culture plates for 24 h. Cells were pre-treated with 0%, 10% and 30% PAM for 3 h before the absolute ATP production rates were measured by Seahorse. The percentage of ATP generated from glycolysis was shown in black, while that from Oxidative Phosphorylation (OXPHOS) was in grey. After being seeded in the Seahorse XF cell culture plates for 24 h, cells were starved with serum free medium for 3 h and treated with or without EGF, 30% PAM, EGF+30% PAM for 30 min. The cells’ metabolic energy levels were tested by Seahorse XFe96 Analyzer. N=3. **d** Effect of 10PAM concentration on the viability of MDA-MB-231 and MDA-MB-468 cells. Cells were plated in 96-well-plates for 24 h. After 3 h starvation in serum free medium, cells were treated with 10 ng/mL EGF, 70% PAM and 10 ng/mL EGF+different PAM doses (100%, 70%, 50%, 30% and 10%) for 12 h, after which the cell viability was tested. **e** Effect of EGF concentration on the viability of MDA-MB-231 and MDA-MB-468 cells. Cells were plated and starved as per (**d**) and treated with 10 ng/mL EGF, 50% PAM and PAM+different doses of EGF (10 ng/mL, 1 ng/mL, 0.1 ng/mL, 0.01 ng/mL) for 12 h, after which the cell viability was tested. N=3. **f** Effect of the order of adding EGF and PAM on the viability of MDA-MB-231 and MDA-MB-468 cells. The cells were plated and starved as per (**a**,**b**), and treated with the 10 ng/mL EGF, 70% PAM, EGF+PAM as the positive controls. ‘EGF activated’ means the EGF added into serum free medium was activated by plasma jet prior to addition to cells. For ‘EGF activated+10 ng/mL EGF’, EGF was added into the ‘EGF activated’ medium again. Other groups of cells were pre-treated with 10 ng/mL EGF for 30 min and then the PAM added, or pre-treated with PAM first and then EGF added. After 12 h treatment, cell viability was tested. N=3. **g** Effect of different growth factors in creating synergies with PAM on the viability of MCF-7, MDA-MB-468, SUM-159PT and MDA-MB-231 cells. Cells were plated in 96-well-plates at 5,000 cells per well for 24 h and pre-treated with serum free DMEM for 3 h. Cells were then treated with serum free DMEM, or 50% PAM, or 50% PAM+EGF, TGFα, TGFβ, FGF1, FGF2, or FGF10 for 12 h, all at 10 ng/mL. N=3. Asterix indicates significance as compared with ‘PAM’.

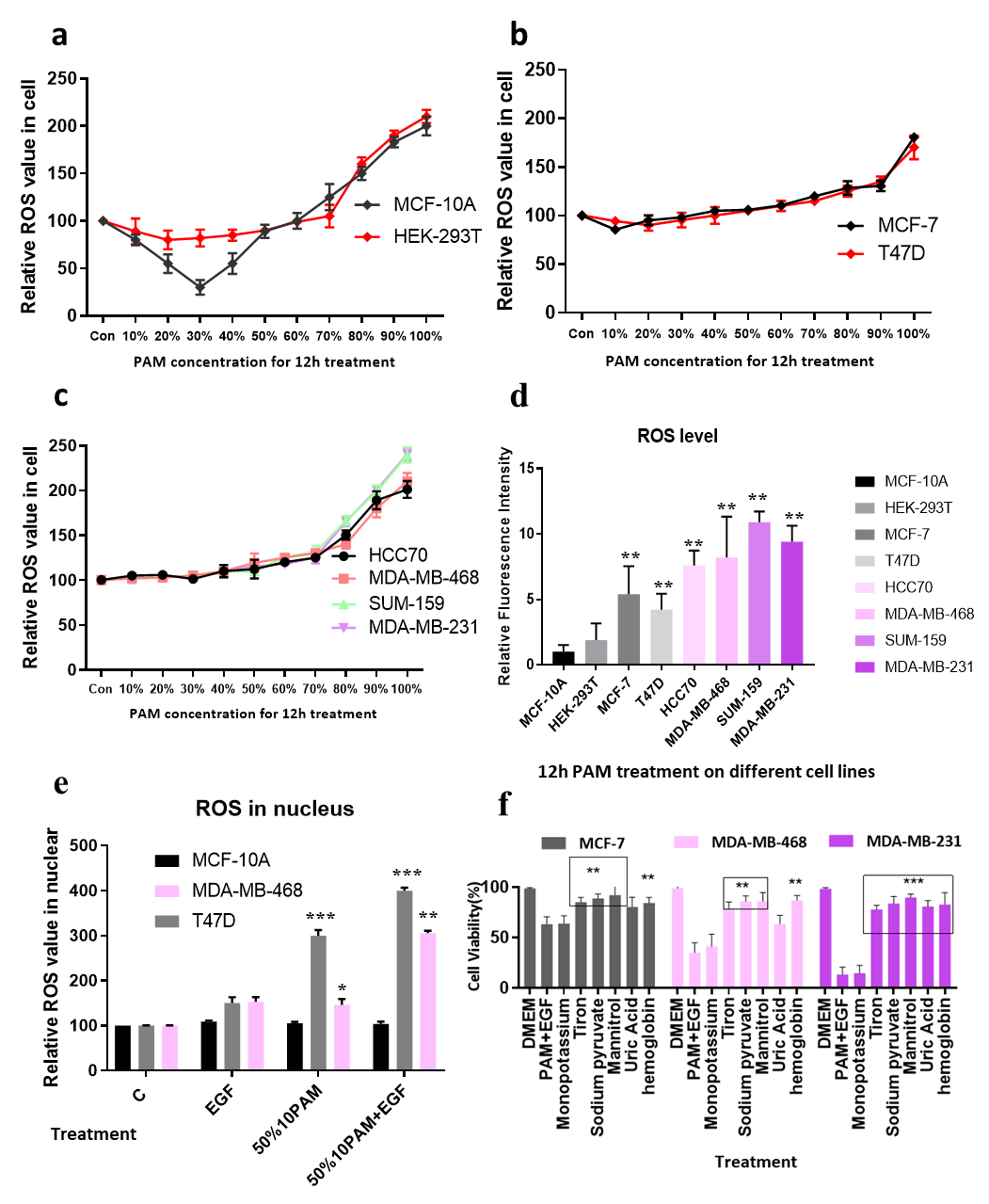

**Supplementary Figure 6. Additional data of Figure 3.** Relative cellular ROS level in response to PAM treatment of different concentrations in (**a**) MCF-10A, HEK-293T; (**b**) MCF-7, T47D; (**c**) HCC70, MDA-MB-468, SUM-159PT and MDA-MB-231 cells. Cells were plated in 96-well-plates at 5,000 cells per well for 24 h, and treated with 0%, 10%, 20%, 30%, 40%, 50%, 60%, 70%, 80%, 90% and 100% of PAM for 12 h. Cells were stained by CellROX® Reagent for 30 min and NucBlue™, images were taken by InCell 6500HS, and ROS levels were calculated by the IN Carta system. **d** The resting ROS level of different breast cancer cell lines. Cells of eight BC cell lines were plated in 96-well-plates at 5,000 cells per well for 24 h and stained with CellROX® Reagent for 30 min. Cells were then stained with NucBlue™ Live Cell Stain and images were taken by InCell 6500HS. Fluorescence intensity was calculated by IN Carta system. N=3. **e** Relative ROS value (%) in the nucleus of MCF-10A, MDA-MB-468 and T47D cells in response to EGF, PAM, EGF+PAM treatments. Cells were treated with or without 100% EGF, 50% 10PAM, or both. **f** Viabilities (% of control; mean +/- SEM; N=3) of MCF-7, MDA-MB-468 and MDA-MB-231 cells pre-treated for 1h with control or scavengers of different ROS components (200 mM mannitol, 100 μM uric acid, 20 mM tiron, 20 μM hemoglobin, 10 mM sodium pyruvate and 1 mM monopotassium were used to quench hydroxyl radical, ozone, superoxide anion, nitric oxide, H_2_O_2_, and e^-^, respectively), and then treated with 50% 10PAM plus 100% EGF.

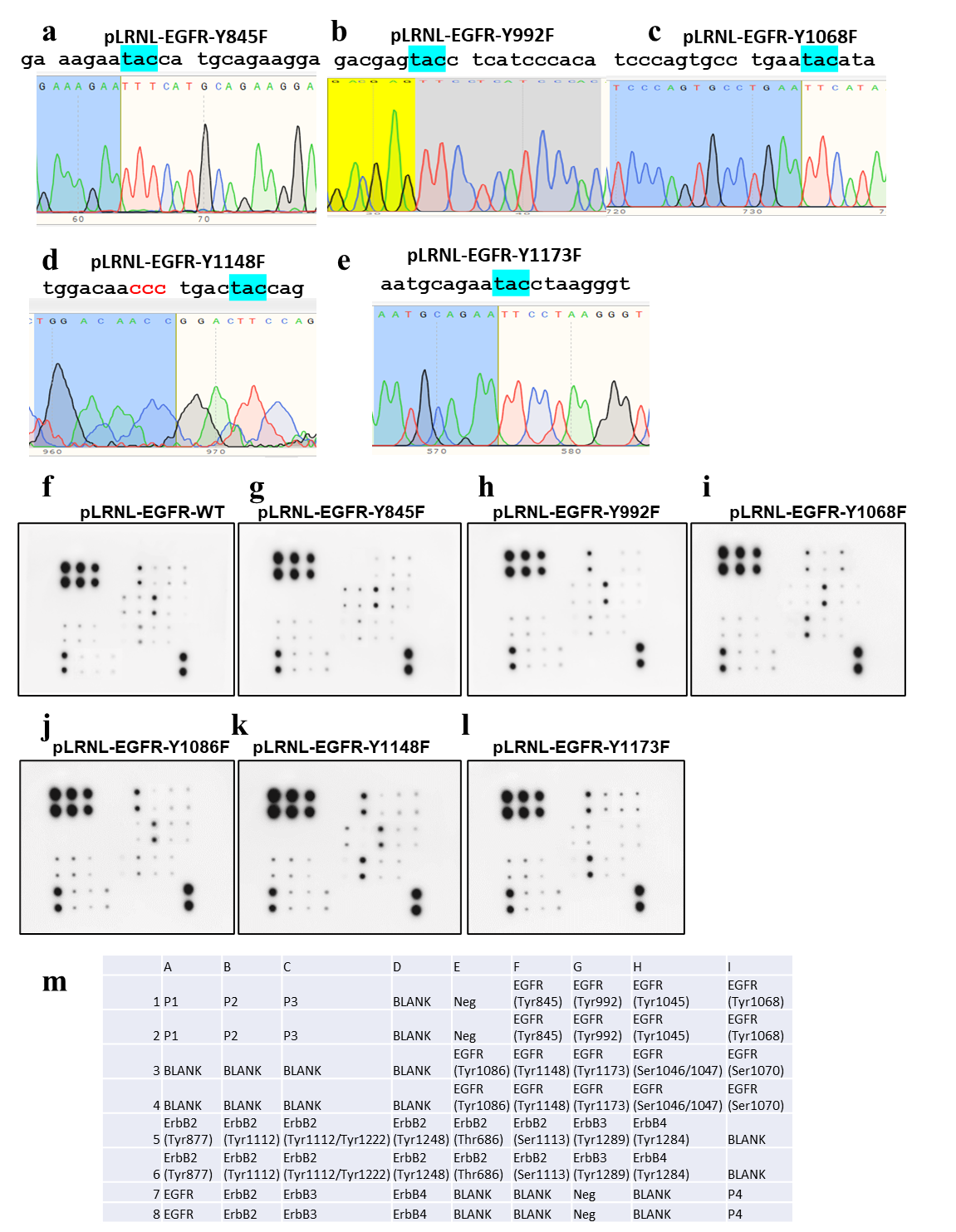

**Supplementary Figure 7. Additional data of Figure 4.** The DNA sequencing results for (**a**) pLRNL-EGFR-Y845F, (**b**) pLRNL-EGFR-Y992F, (**c**) pLRNL-EGFR-Y1068F, (**d**) pLRNL-EGFR-Y1148F, and (**e**) pLRNL-EGFR-Y1173F. The membrane array signal profiles of (**f**) pLRNL-EGFR-WT, (**g**) pLRNL-EGFR-Y845F, (**h**) pLRNL-EGFR-Y992F, (**i**) pLRNL-EGFR-Y1068F, (**j**) pLRNL-EGFR-Y1086F, (**k**) pLRNL-EGFR-Y1148F, (**l**) pLRNL-EGFR-Y1173F in the multi-site EGFR phosphorylation array analysis. EGFR phosphor-site mutated cells were plated at 2$\times$10^7^ cells per T25 flask for 24 h. After 3 h starvation in serum free medium, cells were treated with 10 ng/mL EGF for 30 min, after which protein lysates were collected and analysed as per Material and Methods. (**m**) The phospho-EGFRs antibody data sheet.

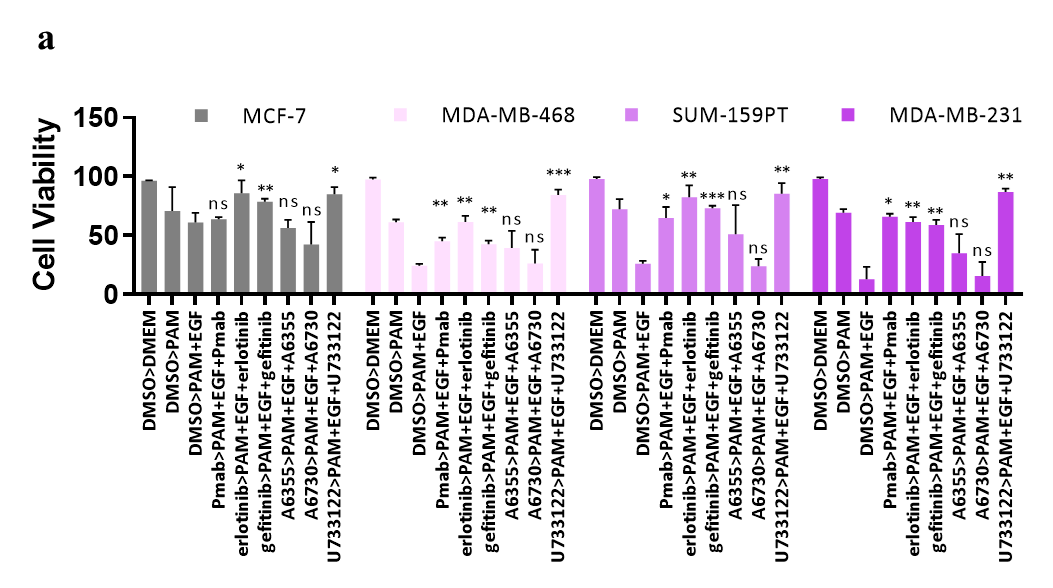

**Supplementary Figure 8. Additional data of Figure 5.** Viabilities of different breast cancer cells pre-treated with different EGFR inhibitors in response to PAM treatment. **a** MCF-7, MDA-MB-468, SUM-159PT and MDA-MB-231 were plated in 96-well-plates at 5,000 cells per well for 24 h. After 3 h serum free medium treatment, cells were pre-treated with DMSO (1 μM), panitumumab (monoclonal antibody of EGFR, Pmab, 50 ng/mL), erlotinib hydrochloride (EGFR tyrosine kinase inhibitor, 1 nM), gefitinib (EGFR tyrosine kinase inhibitor, 17 nM), A6355 (ERK1/2 inhibitor, 5 μM), A6730 (Akt1/2 inhibitor, 210 nM), or U733122 (PLCγ inhibitor, 0.5 μM) in 10% serum DMEM for 1 h. The medium was then changed to serum free DMEM, 50% PAM, or 50% PAM +inhibitors for 12 h. Cell viability was analysed after treatment using the live and dead cell staining kit.

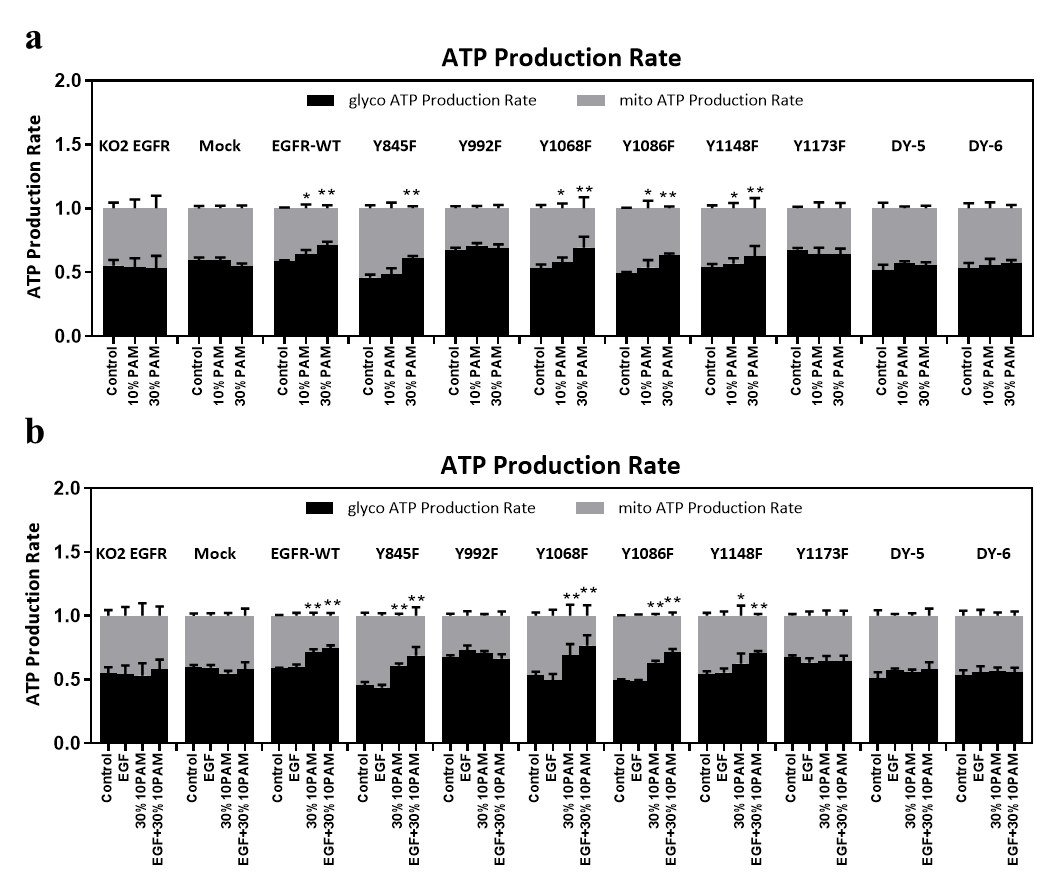

**Supplementary Figure 9.** **Additional data of Figure 6. a** Percentage of ATP production rates of MDA-MB-231 cells transfected without and with different EGFR mutants in response to PAM treatment of different concentrations. MDA-MB-231 KO2 EGFR knockout cells and mock of EGFR transfectants were seeded in Seahorse XF plates at 20,000 per well for 24 h. After 3 h starvation in serum-free medium, cells were treated with or without 10% PAM or 30% PAM for 30 min, after which OCR and ECAR were measured by Seahorse XF Analyzer following the manufacturer's recommendations. The percentage of ATP generated from glycolysis is shown in black, while that from Oxidative Phosphorylation (OXPHOS) is in grey. **b** Percentage of ATP production rates of MDA-MB-231 cells transfected without and with different EGFR mutants in response to EGF, PAM, EGF+PAM treatments. Cells were plated as per (**b**) and were treated with or without EGF, 30% PAM or EGF+30% PAM for 30 min, and the percentages of OCR and ECAR were measured as per (**a**).
